# Modular control of CpG islands by chromatin-opening transcription factors specifies transcription start sites

**DOI:** 10.64898/2026.05.04.722487

**Authors:** Fumiya Moribe, Murat Iskar, Sevi Durdu, Christiane Wirbelauer, Leslie Hoerner, Sébastien A. Smallwood, Lukas Burger, Dirk Schübeler

## Abstract

CpG islands (CGIs) constitute the dominant class of promoters in vertebrate genomes, yet the mechanisms underlying their transcription initiation remain unclear. Using deep learning and acute perturbation of twelve transcription factors (TFs), we show that chromatin-opening TFs are the primary determinants of initiation site selection, thereby governing promoter output. The dominant TF defines a nucleosome-free region and specifies where transcription begins by binding closest to the transcription start site. Loss of this TF redirects transcription initiation to an adjacent secondary TF position, partially buffering gene expression. Promoter variant analyses validate that this modular control of initiation is encoded in *cis* and programmable. This principle extends to initiation at enhancers and TATA-box promoters, where chromatin-opening TFs unmask the TATA box for focused initiation. These findings point to a common sequence grammar for transcription across promoters and enhancers, supporting a minimalist and evolvable architecture of cis-regulatory regions.

## Introduction

CpG islands (CGIs) are a prominent class of gene-proximal regulatory regions in vertebrate genomes and account for the vast majority of RNA polymerase II transcription initiation ^1–3^. Characterized by high CpG density and low DNA methylation, CGIs represent the evolutionarily ancient class of promoters. Across evolution, CGIs maintained high CpG content by remaining hypomethylated in the germline, while the rest of the methylated genome progressively lost CpGs through methylation-dependent C-to-T transitions ^4,5^.

Although their transcriptional regulation remains incompletely understood ^6–9^, the unique chromatin configuration of CGIs, shaped by histone and DNA modifications, has been characterized in greater detail. Such modifications are thought to arise through methylation-dependent recognition of CpGs ^3,7^. Methyl-CpG-binding-domain (MBD) proteins bind methylated CpGs ^10^, while CXXC proteins recognize unmethylated CpGs and mediate deposition of trimethylation at histone H3 lysine 4 (H3K4me3) ^7^. During normal development, only a subset of CGIs becomes DNA-methylated and transcriptionally silenced, enabling allele-specific regulation such as genomic imprinting or X-inactivation ^11–13^. In disease, particularly cancer, aberrant CGI hypermethylation is frequently observed ^14^. By contrast, most CGIs remain unmethylated and exhibit active histone marks and signatures of chromatin accessibility ^3,7^. Consistent with the widespread accessibility of unmethylated CGIs *in vivo* ^15,16^, the intrinsic molecular properties of GC-rich sequences have been shown *in vitro* to repel nucleosome occupancy and recruit chromatin remodelers ^17,18^.

Collectively, these observations have contributed to the view that CGIs are constitutively accessible unless they become DNA-methylated ^6,19^.

Reflective of their evolutionary origin, CGI promoters often drive ancient genes conserved between vertebrates and invertebrates ^8^. These genes typically function in early developmental programs and central metabolic processes. The latter category includes essential genes, commonly referred to as “housekeeping genes”, which are expressed in most cell types, albeit at variable levels associated with diverse proliferative and metabolic states ^8,20^.

CGI promoters display variable and dispersed transcription start sites (TSSs), in contrast to the sharply defined TSSs of TATA-box-containing promoters observed from yeast to humans ^21,22^. The TATA box is a core promoter element bound by the TATA-binding protein (TBP) that supports precise and focused initiation ^23–26^. However, most CGI promoters do not harbor a canonical TATA box, raising the question of how start site selection and transcriptional output are controlled.

Dissecting the regulatory logic of CGIs has been challenging. First, their biased nucleotide composition and the lack of matching CpG-rich genomic sequences without regulatory function limit comparative sequence discovery. Second, CGI activation has been thought to depend on multiple, often redundant regulators, many of which are themselves essential proteins, complicating functional dissection and attribution of individual contributions ^6,27–29^. Lastly, CGI transcription is poorly reconstituted using *in vitro* biochemical assays, which have predominantly relied on TATA-box promoters rather than CGI promoters ^30,31^. Together, these limitations hinder the mechanistic investigation of CGI regulation despite their broad relevance.

Here we combine deep learning with acute TF perturbation to define a cis-regulatory grammar of CGI activation. We show that CGI accessibility is continuously maintained by chromatin-opening TFs, and that promoter output is dominated by the chromatin-opening TF positioned closest to the TSS. This *cis*-encoded modular logic extends to TATA-box promoters and enhancers, supporting a unifying mechanism for transcription initiation across cis-regulatory elements.

## Results

### Deep learning nominates regulatory transcription factors at CpG islands

We first characterized chromatin accessibility as an indicator of regulatory activity using our genome-wide DNase I hypersensitivity map previously generated in mouse embryonic stem cells (mESCs) ^32^. To minimize the confounding effects of DNA methylation, we focused our analysis on unmethylated promoters (82.3%) ^33^. CGIs were generally accessible but individual CGI accessibility varied widely and was higher at transcriptionally active promoters (**Fig. 1a**). This variability suggests that CGIs, often viewed as constitutively accessible ^6,18,34^, may instead be dynamically regulated and become more accessible when actively transcribed. This resembles the context-dependent accessibility at enhancers, a hallmark of cell-type-specific regulatory elements ^35^.

**Fig. 1:**
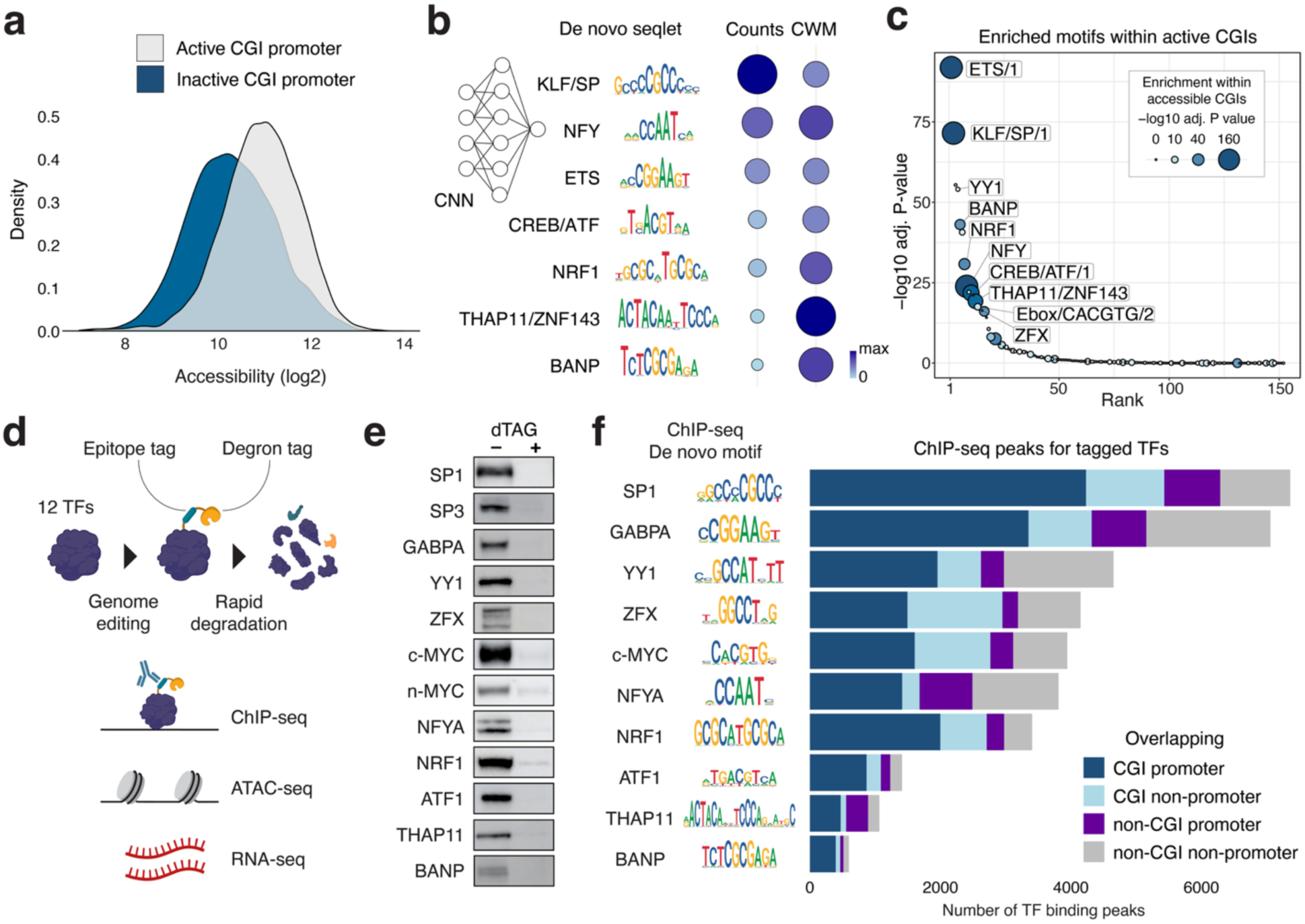
Deep learning model identifies regulatory transcription factors at CpG islands. a, Comparison of chromatin accessibility (log2) measured by DNase-seq between transcriptionally active (n = 7,244) and inactive CGI promoters (n = 2,041) in mESCs. **b,** CNN analysis identifies regulatory sequences (top de novo seqlets) accounting for CGI accessibility in mESCs. Seqlet counts indicate the number of sequences predicted to contribute to higher accessibility, identified by DeepLift combined with TF-MoDISco. Contribution weight matrices (CWM) show nucleotide-level effects across seqlets. **c,** TF motifs enriched in active CGIs, ranked by enrichment (-log10 adj. *P*-value). Circle color and size indicate enrichment in high-versus low-DHS CGIs. **d,** Schematic of the TF engineering strategy enabling epitope-tag-assisted identification of genome-wide binding sites and degron-tag-assisted functional characterization. **e,** Western blot detection of degron-tagged TFs before and after dTAG treatment (500 nM dTAG13 for 4 hours). **f,** ChIP-seq binding profiles of tagged TFs revealing motifs and genomic binding preferences. Left, de novo motifs within the top 1,000 binding peaks. Right, counts of TF binding peaks overlapping CGIs and promoters.

To identify regulatory sequences that account for such variable accessibility, we adapted a convolutional neural network (CNN) model ^36,37^ to predict CGI accessibility from DNA sequences, achieving strong predictive performance (R = 0.72) (**Extended Data Fig. 1a, Methods**). We then applied sequence attribution and motif discovery approaches ^38,39^ to identify motifs driving the predicted accessibility. This analysis revealed a set of motifs recognized by TFs previously implicated in promoter regulation, including KLF/SP, NFY, ETS, CREB/ATF, NRF1, THAP11/ZNF143 and BANP ^32,40–45^ (**Fig. 1b, Extended Data Fig. 1b**). Notably, this set included the BANP motif, consistent with our previous finding that BANP maintains accessibility at a subset of unmethylated CGIs ^45^. As a complementary approach, we performed motif enrichment analysis (monaLisa) ^46^ in transcriptionally active CGIs but not specifically associated with accessibility. This approach recovered motifs similar to those from the CNN model, and additionally revealed motifs for MYC, YY1 and ZFX (**Fig. 1c**), which have been associated with CGI binding ^47–49^. To study the regulatory impact of these CGI-binding TFs (CGI-TFs), we focused on twelve candidates expressed in mESCs (**Fig. 1e**), spanning a wide range of predicted contributions to accessibility.

### CpG islands are selectively targeted by transcription factors

Motif-based predictions of regulatory impact are inherently limited because most TFs occupy only a small fraction of cognate-motif instances in the genome ^50,51^. We therefore sought to define the function of the CGI-TFs at their bona fide binding sites. Stable genetic deletions are not suitable for our candidate TFs because most of them are essential for cell viability based on large-scale loss-of-function screens ^52^. Instead, we used CRISPR-Cas9 to insert a degron and epitope fusion tag at each endogenous TF locus in mESCs, enabling rapid and reversible degradation with the dTAG system ^53^ (**Fig. 1d**). Genome editing was validated by PCR and acute depletion by western blotting for each TF (**Fig. 1e, Extended Data Fig. 1c, Methods**). Tagging endogenous NRF1 was challenging due to the expression of multiple isoforms. We therefore expressed two ectopic degron-tagged isoforms from a chromosomal landing site and subsequently deleted the endogenous *Nrf1* alleles (**Methods**). To account for redundancy among TFs that recognize the same motif ^54,55^, we co-tagged c-MYC/n-MYC and SP1/SP3 as pairs in the same cell lines. Among ∼20 TFs potentially binding the CREB/ATF motif (CRE), we selected ATF1 for tagging based on its high expression in mESCs ^56^. Overall, this effort yielded twelve engineered TFs, representing a broad set of candidates with potential roles in CGI regulation. Following extended degradation times (> 24 h), impaired proliferation or loss of viability was frequently observed, consistent with their essential roles in mESCs.

Next, we determined genomic binding for each TF by ChIP-seq. Using the epitope tag inserted alongside the degron tag, we circumvented the need for ChIP-grade antibodies for each TF and variability in antibody specificity and affinity. This strategy yielded robust binding profiles, enabling systematic analysis of TF occupancy as a function of motif and chromatin state (**Extended Data Fig. 1d**). De novo motif discovery within the top 1,000 peaks recovered the expected motif for each TF (**Fig. 1f**), which closely mirrored the motifs obtained from our CNN approach (**Fig. 1b**). As expected, only a small fraction of the motif instances in the genome were occupied (0.3% for NFYA to 9.6% for SP1, **Extended Data Fig. 1e**), highlighting the importance of experimental validation of binding sites. Binding was nevertheless motif-specific as the cognate motifs were present in the majority of strongly bound peaks in all cases (**Extended Data Fig. 1f**). While binding was highly enriched at CGIs (average 66% of binding peaks), each TF occupied only a limited subset of CGIs (**Fig. 1f, Extended Data Fig. 1g**). Taken together, these results indicate that individual TFs target distinct sets of CGIs in a motif-dependent manner, conferring regulatory specificity in CGI activation.

### CpG island accessibility is driven by transcription factors

To quantify how individual CGI-TFs contribute to chromatin accessibility, we performed acute TF degradation followed by ATAC-seq. Consistent with distinct binding to CGIs (**Fig. 1f**), TF degradation induced selective, factor-specific responses in promoter accessibility. For example, degradation of the CNN-predicted TF GABPA reduced accessibility locally at its bound promoters (**Fig. 2a,b top**). By contrast, degradation of c-MYC/n-MYC, TFs not predicted to increase accessibility, had minimal effects at most bound sites. Across the predicted TFs (**Fig. 1b**), their degradation frequently decreased accessibility at bound promoters (**Fig. 2b top**), supporting the model’s prediction. The magnitude of accessibility change correlated with binding strength, suggesting a direct, binding-site-specific response.

**Fig. 2:**
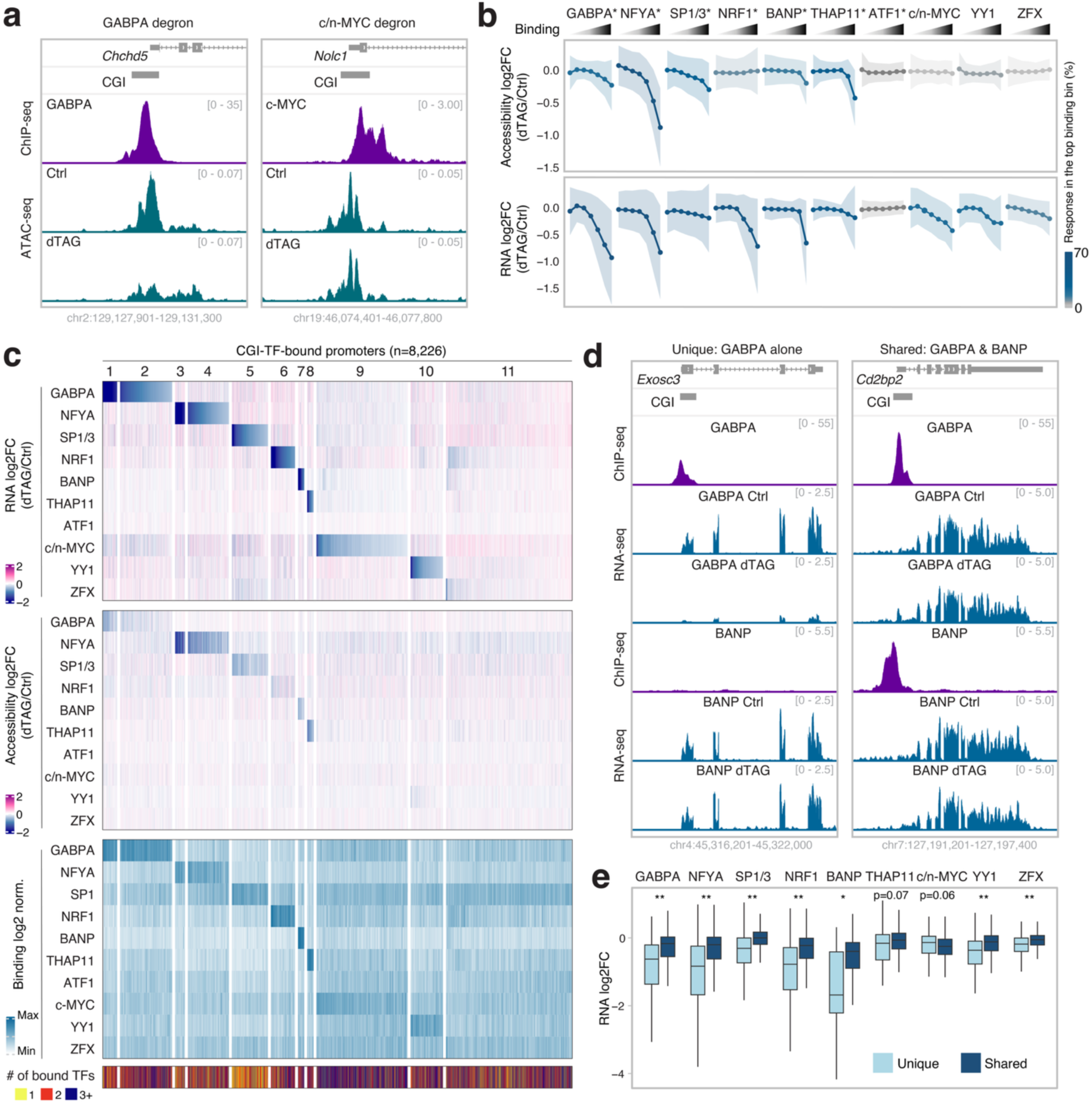
Regulatory specificity of transcription factors at CpG islands. a, Representative loci illustrating TF binding (ChIP-seq) and chromatin accessibility changes (ATAC-seq) upon TF degradation (500 nM dTAG13 for 4 hours). **b,** Changes in chromatin accessibility and transcription upon TF degradation as a function of binding strength at promoters. Top: Mean ATAC-seq fold change (log2, dTAG/Ctrl) at active promoters (n = 10,905, CGI promoters; 76%) grouped by binding strength of the respective TF (ChIP-seq enrichment ranked by 20-20-20-15-10-10-5%, from weakest to strongest). Shaded areas represent standard deviation, and line colors represent fraction of significant response in the top bin (FDR < 0.05 & |log2 fold change| > 0.5). Asterisks indicate CNN-nominated TFs. Bottom: same analysis using RNA-seq. **c,** Genome-wide effects of TF degradation on transcription and accessibility. Columns represent individual promoters bound by at least one of the TFs (n = 8,226, CGI promoters; 90%). Panels display fold change of RNA-seq, ATAC-seq, and scaled enrichment of ChIP-seq (log2). Bottom bar indicates number of bound TFs. Promoters are *k*-means clustered by RNA-seq change and sorted by downregulation strength. **d,** Representative loci comparing transcriptional changes upon GABPA or BANP degradation at promoters either bound by GABPA (Unique) or by both GABPA and BANP (Shared). **e,** Comparison of transcriptional responses upon TF degradation at promoters bound by a single TF (Unique) or multiple TFs (Shared). Promoter counts (Unique/Shared) are GABPA: 384/3,915, NFYA: 218/1,969, SP1/3: 700/4,162, NRF1: 154/2,094, BANP: 25/450, THAP11: 62/790, c/n-MYC: 102/1,857, ZFX: 106/1,548. Wilcoxon rank-sum test: \**P*-value < 1.0e-3, \*\**P*-value < 1.0e-5.

Among tested candidates, NRF1 degradation had only a limited impact (**Fig. 2b top**), despite previous reports that NRF1 can open chromatin ^32,57,58^. However, the degron tagging itself already reduced accessibility at NRF1 binding sites prior to degradation (**Extended Data Fig. 2a,b**). This pre-existing reduction likely reflects compromised activity of the tagged protein and/or incomplete isoform representation (**Methods**), limiting our ability to detect the full effect of NRF1. A second outlier was ATF1: its degradation caused no detectable accessibility change, possibly due to redundancy among other CRE binders that open chromatin ^43,56,58,59^.

Overall, CNN nomination was broadly consistent with perturbation data, with exceptions likely reflecting factor-specific limitations from the tagging strategy or redundant binders. Collectively, we classified the CGI-TFs into chromatin-opening (GABPA, NFYA, SP1/3, NRF1, BANP and THAP11), and non-chromatin-opening (c-MYC/n-MYC, YY1, and ZFX).

Although the accessibility loss upon TF degradation demonstrates that chromatin-opening TFs are required for maintaining open chromatin, it does not address whether they are sufficient for establishing it. To test this, we degraded GABPA or BANP for 4 hours, washed out dTAG and allowed the TF levels to recover during the subsequent 16 hours. TF recovery restored accessibility to baseline levels at their target sites, as determined by ATAC-seq (**Extended Data Fig. 2c,d**). Thus, these TFs are sufficient to re-establish accessible chromatin at their target CGIs, suggesting that chromatin-opening at CGIs is a continuous, TF-driven process. Together, these results indicate that open chromatin at CGIs is not a default state but is dynamically regulated by a defined set of sequence-specific TFs.

### CpG island activation is transcription-factor-specific

Having shown that a distinct set of CGI-TFs can open chromatin at CGIs, we next investigated how these changes translate into transcriptional output. We performed total RNA-seq eight hours after inducing degradation, a time point chosen to capture primary transcriptional changes while minimizing subsequent secondary effects ^45^. Across perturbations, CGI-TF degradation resulted in differential expression of 6,935 genes, revealing regulatory effects for nearly two-thirds of all expressed genes (**Extended Data Fig. 2e**). In accordance with their roles as activators, degradation of candidate TFs resulted primarily in downregulation of genes associated with bound promoters (**Fig. 2b bottom, Extended Data Fig. 2f**). This pronounced enrichment of responses at bound genes indicates that these genes are primary targets, supporting the specificity of acute degradation. Consistently, the magnitude of the transcriptional response scaled with TF binding strength at promoters (**Fig. 2b bottom, Extended Data Fig. 2f**). Notably, transcriptional downregulation was observed for both chromatin-opening and non-chromatin-opening TFs. In contrast, ATF1 degradation did not lead to detectable transcriptional changes, in line with the absence of accessibility effects, further supporting the redundancy of CRE binders. In support of this, the individual degradation of c-MYC or SP1 alone had minor changes in gene expression, while the combinatorial degradation of c-MYC/n-MYC or SP1/SP3 showed stronger changes, likely reflecting redundancy of factors acting on the same motifs (**Extended Data Fig. 2g**).

We next examined whether changes in accessibility and transcription upon individual TF degradation are linked. For chromatin-opening TFs, accessibility and expression changes were largely concordant (**Fig. 2b**), suggesting that the abilities to open chromatin and to drive transcription are generally coupled for these factors. By contrast, this was not the case for non-chromatin-opening TFs, where we observed downregulation of bound genes despite largely unchanged accessibility (**Fig. 2b**). These results suggest that the ability to open chromatin is not necessarily required for a TF’s ability to activate transcription, pointing to potential mechanistic differences between these TF classes.

To gain insight into the specificity of transcriptional effects, we clustered bound promoters based on their transcriptional responses and compared them to accessibility changes and TF binding (**Fig. 2c**). This comparison illustrated the coupled changes in transcription and accessibility upon degradation of chromatin-opening TFs and furthermore revealed highly TF-specific transcriptional responses (e.g. cluster 1-8 in **Fig. 2c**). This is exemplified by the *Exosc3* promoter bound uniquely by GABPA, whose degradation resulted in strong transcriptional reduction (**Fig. 2d left**). Strikingly, we found this selective response to a single TF even at co-bound promoters.

Specifically, 43% of promoters bound uniquely by one TF responded transcriptionally to degradation of that TF. For the promoters bound by multiple TFs, 42% responded to degradation of only one of the binding TFs, whereas only 10% responded to more than one TF degradation (**Extended Data Fig. 2h,i**). Thus, CGI promoters, while typically associated with broadly active genes, can nevertheless rely predominantly on a single activator. This observation challenges the view that CGI transcription is driven by additive activity of multiple TFs, as would be predicted from dense TF occupancy ^6,28,29^.

Despite such predominant single-TF effects, approximately one-third of bound genes showed no significant transcriptional responses to any TF degradation (e.g. cluster 11 in **Fig. 2c**). Even among strongly bound genes, not all of them were downregulated following degradation of the corresponding TF (**Extended Data Fig. 2f**). This is illustrated at the *Cd2bp2* promoter co-bound by GABPA and BANP, where degradation of neither factor caused a strong transcriptional change (**Fig. 2d right**). Overall, CGI promoters co-bound by multiple TFs exhibited weaker transcriptional responses than those bound uniquely by a single TF (**Fig. 2e**). In contrast, the non-chromatin-opening TF MYC was a notable exception, showing comparable transcriptional effects at both unique and shared promoters despite higher baseline activity of shared promoters (**Fig. 2e, Extended Data Fig. 2j**). This is consistent with previous reports that MYC globally amplifies gene expression, potentially by promoting transcription elongation ^60,61^.

Taken together, these results show that individual TFs activate gene expression at CGIs while only a subset of them can open chromatin. This systematic characterization reveals high TF specificity and diverse promoter responses. These findings challenge a model that CGIs are intrinsically transcriptionally permissive and instead suggest that transcription at CGIs is a highly regulated process.

### Modular control of transcription initiation by chromatin-opening transcription factors

A hallmark of CGIs is their variable usage of transcription start sites (TSSs) ^21^. We therefore asked whether TF specificity and promoter responses are reflected in TSS usage. To map TSSs genome-wide at base-pair resolution, we captured nascent 5’-capped RNAs using NET-CAGE ^62^ (**Fig. 3a**). Applying this method to mESCs yielded highly reproducible data with 95% of reads overlapping FANTOM-annotated TSSs ^20^ (**Extended Data Fig. 3a**). Detected TSSs were enriched at promoters and enhancers (**Extended Data Fig. 3b,c**). CGI promoters often displayed broad TSS profiles manifesting as a cluster of TSS peaks ^21^ (**Fig. 3a**). We thus assigned a single dominant TSS per active promoter based on signal strength (n = 12,631) ^63^ (**Fig. 3a, Methods**).

**Fig. 3:**
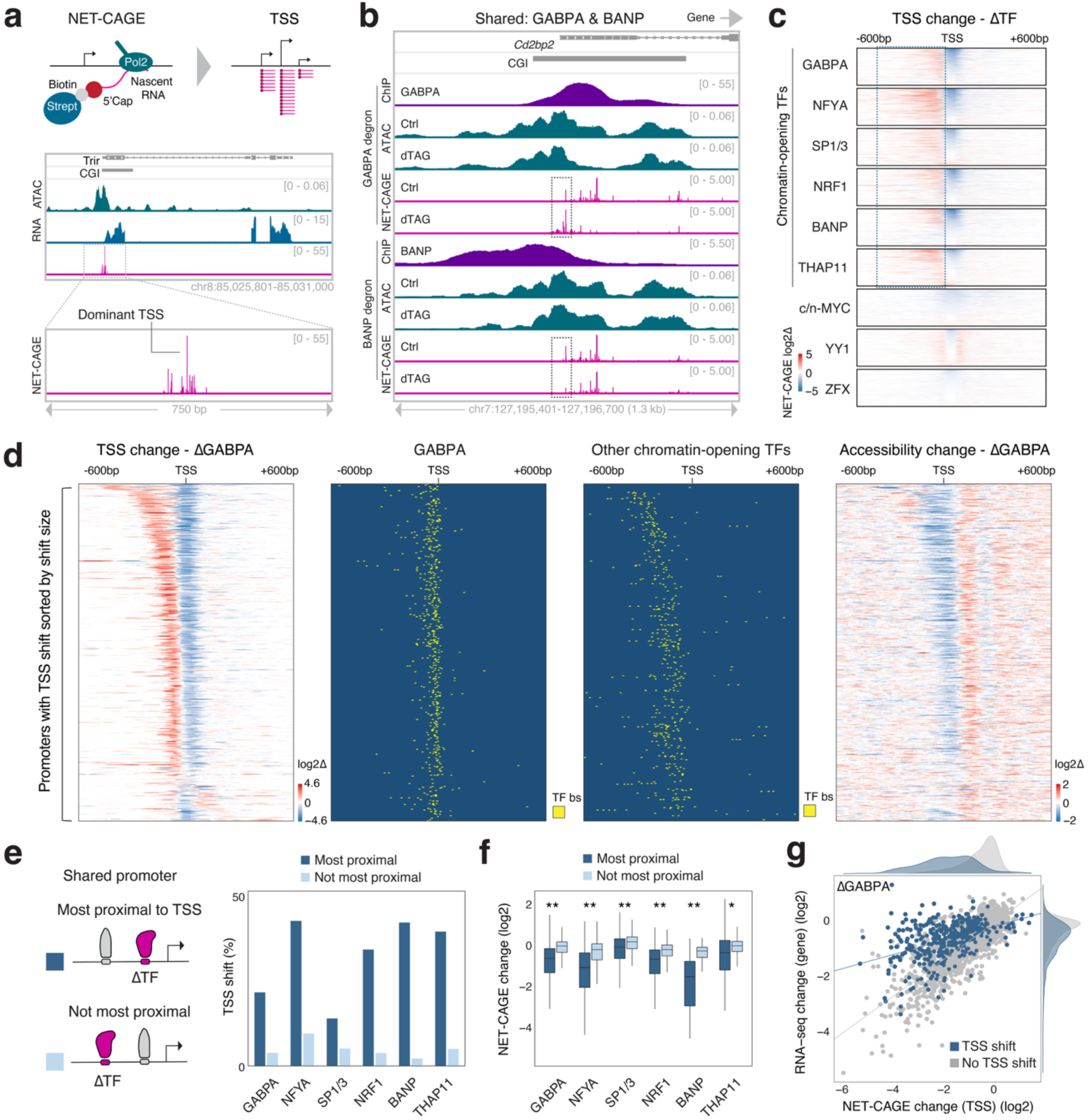
Chromatin-opening transcription factors control transcription initiation and output. a, Schematic of NET-CAGE approach to map TSSs and representative locus illustrating the resulting spatial resolution and a typical TSS profile at CGIs. **b,** Representative locus (*Cd2bp2*) showing a TSS shift upon GABPA degradation. Shown are ChIP-seq, ATAC-seq, and NET-CAGE signal in control and dTAG conditions for GABPA or BANP degron cells. (500 nM dTAG13 for 4 hours). The dotted box marks TSS changes. **c,** NET-CAGE signal changes (log2) at the most strongly bound promoters (top 500) by each TF following its degradation (4,500 promoters in total), ranked by NET-CAGE log2 fold change. Highlighted regions show changes in the alternative TSS signal. **d,** TSS shifts at promoters upon GABPA degradation (n = 417). Panels show NET-CAGE log2 fold change, GABPA bound motifs, bound motifs of other chromatin-opening TFs (NFYA, SP1, NRF1, BANP, THAP11), and ATAC-seq log2 fold change, at promoters ranked by the magnitude of the TSS shift. **e,** Frequency of TSS shifts depends on promoter type. Promoters bound by multiple chromatin-opening TFs are compared, and “Most proximal” indicates that the degraded TF binds closest upstream of the TSS among chromatin-opening TFs. Promoter counts (most proximal/not most proximal) are GABPA: 1,016/989, NFYA: 537/845, SP1/3: 1,006/1246, NRF1: 621/757, BANP: 137/190, THAP11: 175/434. **f,** Similar to **(e)** but showing NET-CAGE fold change (log2) following individual TF degradation. Wilcoxon rank-sum test: \**P*-value < 1.0e-3, \*\**P*-value < 1.0e-5. **g,** Impact of TSS shifts on TSS signal (NET-CAGE log2 fold change) and gene expression (RNA-seq log2 fold change) upon GABPA degradation. Lines indicate linear regression of each group.

Promoters containing a canonical TATA box (-35 bp to-21 relative to the TSS) exhibited sharply defined initiation sites (<10 bp), whereas TATA-less promoters showed broader TSS distributions (∼50 bp), as previously reported ^22^ (**Extended Data Fig. 3d,e**).

Aligning TF binding sites relative to promoter TSSs revealed that chromatin-opening TFs were positioned immediately upstream of dominant TSSs, colocalizing with accessible chromatin (**Extended Data Fig. 3f**). In contrast, binding sites for c-MYC, YY1 and ZFX, whose degradation did not alter accessibility at the majority of bound promoters, were preferentially enriched downstream of the dominant TSSs (**Extended Data Fig. 3f**).

Next, we asked how individual TFs affect promoter transcriptional output and TSS choice by performing NET-CAGE following acute degradation. The resulting datasets identified 9,634 differentially expressed TSSs at promoters, indicating that over three-quarters of all expressed genes responded to at least one of the TF perturbations (**Extended Data Fig. 3g**). Initial binding strength strongly correlated with the magnitude of initiation changes upon degradation (**Extended Data Fig. 3h**), mirroring the effects measured by RNA-seq (**Fig. 2b**). To assess these changes in more detail, we analyzed TSS peak strength and distribution after each TF degradation. Strikingly, reduction of the dominant TSS peak frequently coincided with the upregulation of an initially weaker, alternative upstream TSS. This is exemplified at the *Cd2bp2* promoter, co-bound by GABPA and BANP (**Fig. 3b**), which showed only a minor RNA-seq change upon degradation of either TF (**Fig. 2d**). At this locus, GABPA degradation induced a reduction of the dominant TSS signal accompanied by a marked increase in an alternative peak proximal to the upstream BANP site. Conversely, BANP degradation reduced this secondary TSS signal (**Fig. 3b**). Such TSS shifts towards an upstream (5’) position were observed at over 1,800 promoters, almost exclusively upon degradation of chromatin-opening TFs as non-chromatin-opening TFs accounted for only 4.2% of these shifts (**Fig. 3c, Extended Data Fig. 3i**). YY1 degradation induced TSS shifts to downstream positions (typically +1 to +50 bp) at a small fraction of bound promoters (**Extended Data Fig. 3i**).

The coincidence of a TSS shift with an upstream BANP binding site at the *Cd2bp2* promoter (**Fig. 3b**) prompted us to ask how binding positions of CGI-TFs relate to TSS shifts. For each chromatin-opening TF, we analyzed promoters showing TSS shifts upon its degradation and mapped binding sites for other chromatin-opening TFs at these promoters. Strikingly, upstream-shifted, alternative TSSs were enriched specifically at binding sites for other chromatin-opening TFs (**Fig. 3d, Extended Data Fig. 4a**), suggesting that initiation relocated to adjacent chromatin-opening TFs. This pattern was conserved across all chromatin-opening TFs tested (**Extended Data Fig. 4b-f**). At promoters co-bound by multiple chromatin-opening TFs, TSS shifts occurred preferentially when the degraded TF resided most proximal to the dominant TSS (**Fig. 3e, Extended Data Fig. 4g**). Consistently, degradation of this TSS-proximal TF had stronger effects on initiation output (**Fig. 3f**). This positional effect could explain why many promoters, despite being bound by multiple TFs, responded primarily to degradation of a single factor (**Extended Data Fig. 2i**). At the same time, however, promoters undergoing a TSS switch had a smaller decrease in total RNA-seq signal than a reduction at the initially dominant TSS (**Fig. 3g, Extended Data Fig. 4h**). This suggests that TSS shifts can partially buffer the impact of TF ablation on gene expression.

Given that transcription initiation switches between binding sites for chromatin-opening TFs, we asked whether TSS changes are coupled to accessibility responses. We compared NET-CAGE and ATAC-seq changes across promoters that showed TSS shifts upon TF degradation (**Fig. 3d, Extended Data Fig. 4b-f**). This revealed a spatially confined loss of accessibility around downregulated TSSs, directly linking TF-dependent TSS selection to local chromatin opening and suggesting that local accessibility can be a prerequisite for transcription initiation.

Together, these results identify TF binding as a primary determinant of TSS selection at CGI promoters, which appears linked to chromatin-opening activity. At most CGI promoters, a single chromatin-opening TF positioned closest to the TSS dominates promoter output, consistent with TF-specific transcriptional responses. At a subset of promoters co-bound by multiple chromatin-opening TFs, loss of the dominant TF induces a switch to an alternative TSS, providing partial but measurable buffering of gene expression. This hierarchical yet locally redundant architecture offers a mechanistic framework for how CGI promoters can be both selectively dependent on a single chromatin-opening TF as well as resilient to perturbation in specific configurations. These findings suggest that CGI promoters – often controlling essential genes – can exhibit regulatory flexibility, where binding by multiple chromatin-opening TFs can buffer against strong expression changes.

### Gene-proximal transcription factors specify initiation sites

The observation that acute TF depletion in *trans* induced TSS shifts towards upstream TF positions supports a model in which the most gene-proximal chromatin-opening TF specifies the dominant initiation site. If true, this model would predict that TSS selection is promoter-autonomous and can be forcibly redirected by repositioning TF motifs. To test these predictions, we engineered variants of endogenous CGI promoters harboring defined motif insertions and deletions. Each variant was coupled to a luciferase coding sequence and integrated into a fixed chromosomal landing site by recombination-mediated cassette exchange (RMCE) ^64^ (**Fig. 4a, Extended Data Fig. 5a, Methods**). To quantify how motif perturbations alter TSS choice at the engineered locus, we developed locus-specific CAGE. This method selectively captured transcripts from the integrated promoters, and promoter-unique barcodes encoded in the transcripts enabled assignment of CAGE reads to their corresponding promoter variants (**Extended Data Fig. 5a, Methods**). In parallel, we combined locus-specific CAGE with single-molecule footprinting (SMF) to precisely map nucleosome occupancy and TF binding at the promoters ^65^, directly linking local chromatin organization with transcription initiation (**Fig. 4a**).

**Fig. 4:**
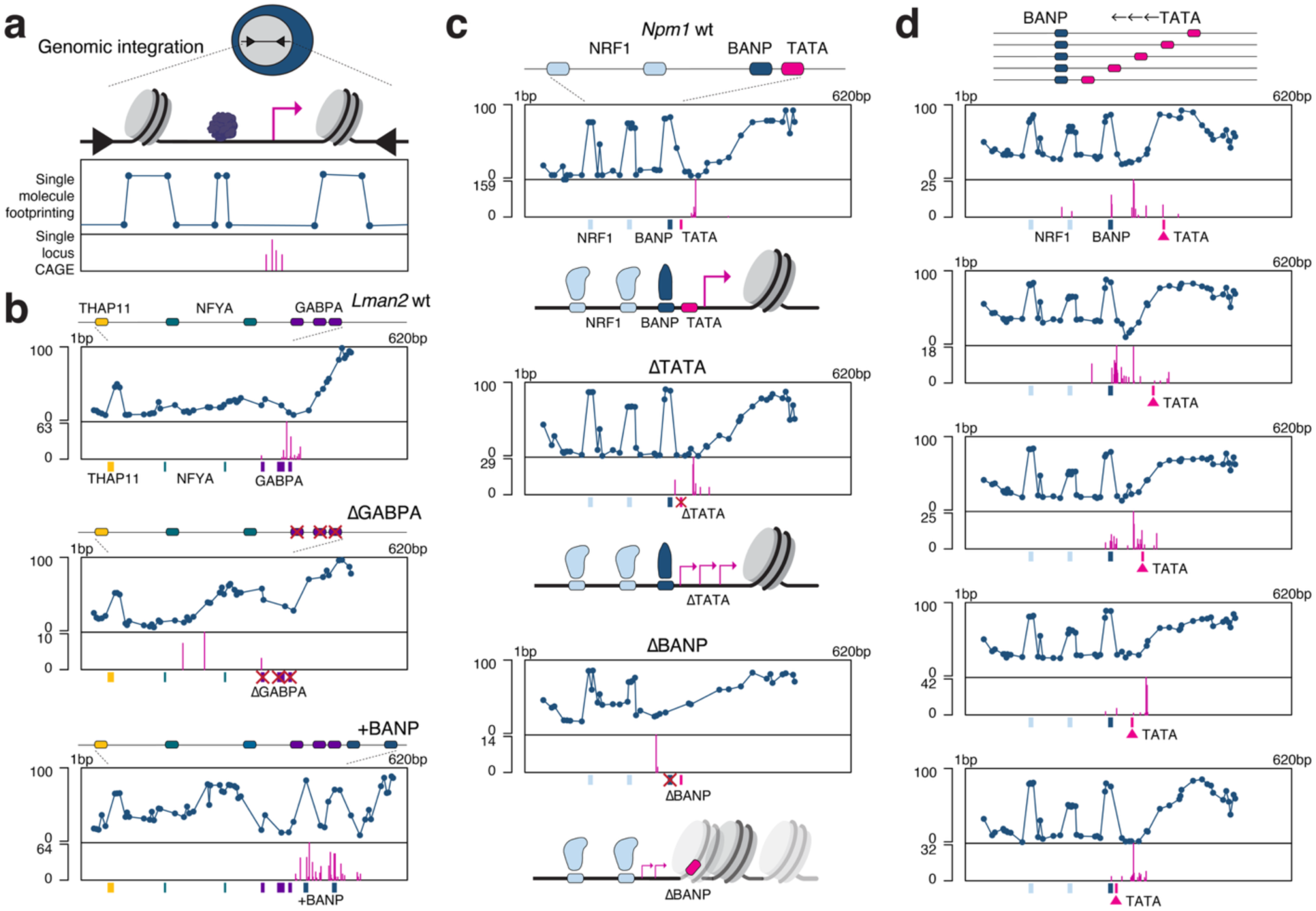
Control of nucleosome positioning and initiation sites by transcription factors. a, Schematic of locus-specific CAGE combined with GpC single-molecule footprinting (SMF) to monitor TSS usage and nucleosome occupancy. **b,** Analysis of the *Lman2* promoter bound by GABPA, NFYA and THAP11. Upper panels display SMF signal (dotted line, 100 - methylation %) and lower panels show locus-specific CAGE TSS signal (vertical stripes). Top: WT promoter sequence recapitulates the endogenous signal. Middle: mutation of GABPA motifs increases nucleosome occupancy (broad SMF peaks) over the mutated motifs and initial position of TSSs, resulting in a TSS shift. Bottom: insertion of BANP motifs induces nucleosome depletion and TSS relocation (BANP occupancy also generates TF-specific narrow footprinting peaks ^65^). **c,** Analysis of a TATA-box-containing CGI promoter (*Npm1*). Top: WT. Middle: TATA box mutation results in a dispersed TSS signal. Bottom: BANP motif mutation leads to increased nucleosome occupancy over the TATA box and a TSS shift towards NRF1 binding sites. Each schematic illustrates chromatin configurations of the respective promoter. **d,** Positional effects of TATA box insertion downstream of the BANP motif. TATA boxes are placed at varying distances from the BANP motif (20 bp intervals from 5’ to 3’).

As proof-of-principle, we first quantified TSS usage of wild-type promoter sequences using locus-specific CAGE and SMF. The integrated *Lman2* promoter recapitulated the initiation pattern of the endogenous locus, with predominant signal at the gene-proximal GABPA motifs (**Fig. 4b, Extended Data Fig. 5b**), suggesting that TSS positioning is promoter-autonomous. Moreover, mutating GABPA motifs caused a TSS shift from the GABPA-defined position towards upstream NFYA motifs, concomitant with loss of SMF accessibility over the original TSS at the GABPA sites (**Fig. 4b**). This mirrored the response at the endogenous promoter upon GABPA degradation (**Extended Data Fig. 5b**). We next asked whether introducing an even more gene-proximal binding site for a chromatin-opening TF would reposition the TSS in the opposite direction. Indeed, inserting BANP motifs closer to the gene redirected the dominant TSS downstream (**Fig. 4b**), supporting the model that the most gene-proximal chromatin-opening TF defines the dominant start site. Concomitantly, the BANP motif insertion displaced the +1 nucleosome downstream, creating a nucleosome-free region at the new TSS.

We next examined the *Rps21* promoter, which contains motifs for both the chromatin-opening TF GABPA and the non-chromatin-opening TF MYC. Consistent with TF degradation at the endogenous locus, deletion of either the GABPA or MYC motifs reduced TSS activity, with a stronger effect observed upon GABPA loss (**Extended Data Fig. 5c, d**). Importantly, GABPA ablation led to full nucleosome occupancy over the initial TSS position and near-complete loss of initiation. By contrast, deleting the MYC motif did not alter nucleosome occupancy and only partially reduced TSS activity.

Together, these results reinforce a model in which the most gene-proximal chromatin-opening TF establishes accessible chromatin and specifies TSS positions, while additional TFs enhance transcriptional output without dictating nucleosome organization.

### Transcription-factors license TATA box

Having defined how chromatin-opening TFs position TSSs at CGI promoters, we next asked how this TF-driven mechanism interfaces with the canonical core promoter element, the TATA box. Although TATA boxes are not common at mammalian promoters including CGIs, TATA-box promoters overlap CGIs at a frequency comparable to that of TATA-less promoters (**Extended Data Fig. 5e**). TATA-containing CGI promoters include strong promoters widely used in transgenic expression constructs, such as those driving *Gapdh*, *Ef1a* and *Actb*. The *Npm1* promoter exemplifies this architecture: it displays a prominent NET-CAGE peak downstream of the TATA box, and is co-bound by BANP and NRF1, with BANP closest to the TATA box (**Extended Data Fig. 5f**). When inserted into the chromosomal landing site, the *Npm1* promoter recapitulated sharp, focused initiation at the TATA box observed at the endogenous locus (**Fig. 4c top**).

Scrambling the TATA box retained a similarly accessible chromatin state but produced a more dispersed initiation pattern (**Fig. 4c middle**), resembling that of TATA-less CGI promoters. Reciprocally, introducing a TATA box into the otherwise TATA-less CGI promoter of *Lman2* converted its dispersed initiation into a sharp peak (**Extended Data Fig. 5g**). Combined, these data indicate that CGI promoters possess the plasticity to support both sharp and broad initiation depending on the presence of a TATA box.

Next, we hypothesized that gene-proximal binding of a chromatin-opening TF is required to render the TATA box accessible and functional. To test this model, we mutated the BANP motif adjacent to the TATA box at the *Npm1* promoter. This mutation strongly reduced TSS signal at the TATA box and induced a pronounced TSS shift towards an upstream NRF1 site (**Fig. 4c bottom**). SMF revealed that BANP motif mutation also repositioned the +1 nucleosome over the TATA box. Consistently, BANP degradation at the endogenous *Npm1* promoter reduced the dominant TSS peak and increased an upstream initiation proximal to the NRF1 site (**Extended Data Fig. 5f**). Combined, these results support a model in which a chromatin-opening TF in the vicinity of the TATA box is necessary for its productive usage. This model predicts positional constraints of TATA box usage, which we tested systematically by placing the TATA box at variable distances downstream of the BANP motif. The resulting TSS/chromatin profiles revealed that efficient usage of the TATA box requires that it resides within the nucleosome-free region created by BANP (**Fig. 4d**). More distal positions failed to support focused initiation and instead showed nucleosome occupancy over the TATA box. Altogether, the systematic dissection of the *Npm1* promoter suggests that chromatin-opening TFs unmask the TATA box from nucleosome occlusion and license TATA-box usage in their vicinity.

We next tested whether this principle generalizes genome-wide. Many TATA-box promoters were bound by chromatin-opening TFs (**Extended Data Fig. 5h**), with binding sites located immediately upstream of the TATA box (**Extended Data Fig. 5i**). At these promoters, indeed, degradation of chromatin-opening TFs frequently led to transcriptional downregulation and TSS shifts, as predicted (**Extended Data Fig. 5j,k**).

Taken together, *cis* and *trans* perturbations demonstrate that gene-proximal chromatin-opening TFs create nucleosome-free regions within CGIs and specify TSSs independently of the presence of a TATA box. These findings suggest that chromatin-opening TFs can be sufficient to license transcription initiation at CGIs.

### Chromatin-opening transcription factors drive enhancer RNA initiation

Our finding that chromatin-opening TFs control TSS selection at promoters raises the possibility that they also govern transcription initiation at distal regulatory elements such as enhancers. Similar to promoters, active enhancers are characterized by TF occupancy and accessible chromatin and produce enhancer RNAs (eRNAs) ^66–70^. eRNAs are short non-coding transcripts (∼300 nt), often produced bidirectionally, and have been implicated in distal gene regulation ^71,72^. Despite the short half-life of eRNAs, NET-CAGE detected TSSs at candidate transcribed enhancers (n = 2,640) (**Fig. 5a, Methods**). Relative to promoter TSSs, enhancer TSSs showed lower expression and reduced accessibility (**Extended Data Fig. 6a,b**). Only 17% of active enhancers overlapped CGIs ^73,74^, resulting in lower CGI-TF occupancy than at promoters ^29^ (**Extended Data Fig. 6c,d**). Upon TF degradation, individual candidate enhancers showed pronounced reductions in NET-CAGE signal (**Fig. 5b**). Genome-wide downregulation of enhancer TSS signal scaled with TF binding strength (**Fig. 5c, Extended Data Fig. 6e**), similar to what we observed at promoters (**Fig. 2b**). These data suggest that chromatin-opening TFs are required for transcription initiation at a subset of distal non-promoter sites.

**Fig. 5:**
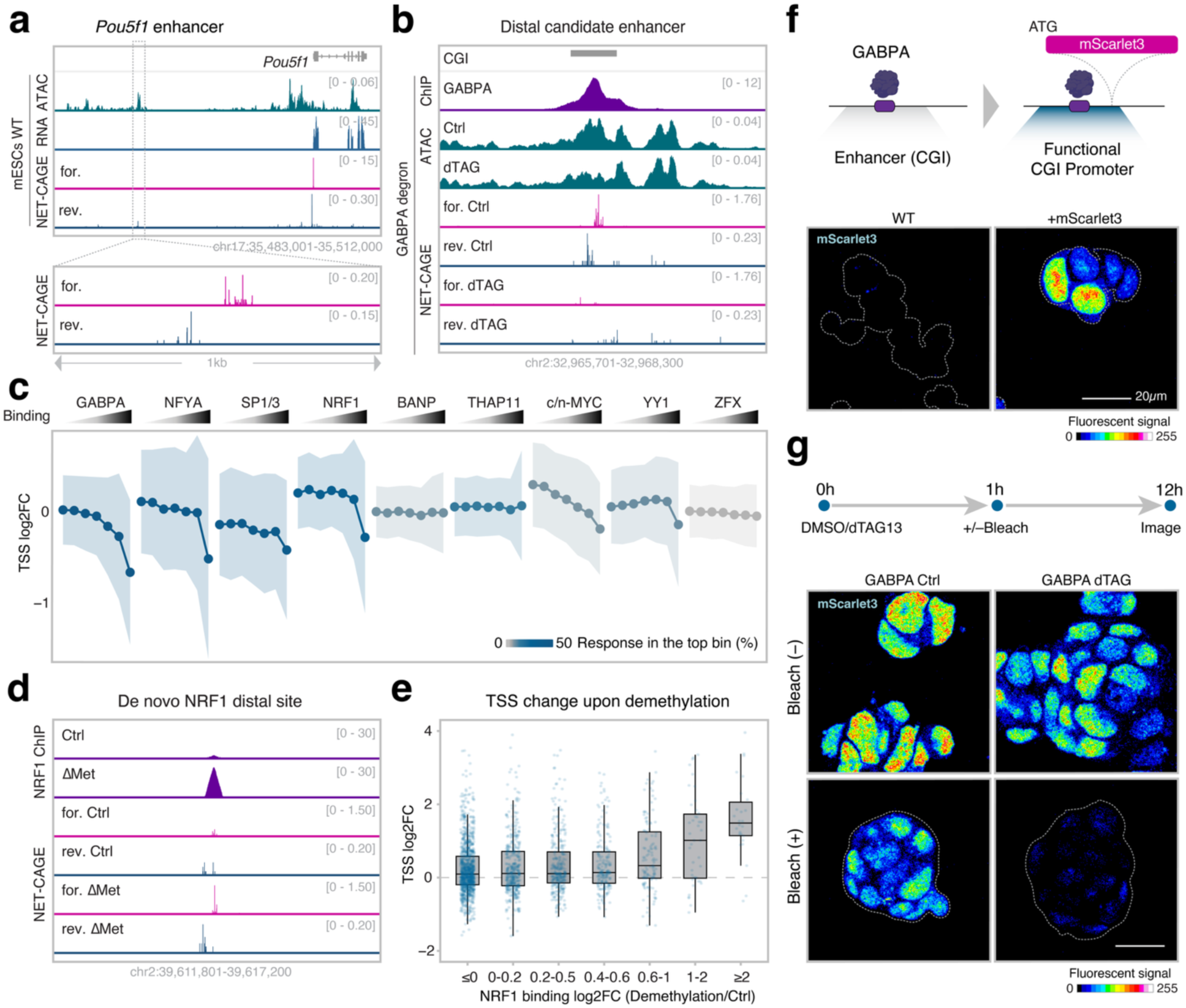
Transcription factors drive initiation independently of coding sequences. a, Representative locus (*Pou5f1*) showing transcription initiation at promoter and enhancer, measured by ATAC-seq, RNA-seq, and NET-CAGE. **b,** Representative candidate enhancer illustrating GABPA-dependent eRNA production. ChIP-seq, ATAC-seq and NET-CAGE signal for control and GABPA-degraded conditions (500 nM dTAG13 for 4 hours) are shown. **c,** Changes in NET-CAGE signal upon TF degradation as a function of binding strength at candidate enhancers (n = 2,640). Mean NET-CAGE fold change (log2, dTAG/Ctrl) at active enhancers grouped by binding strength of the respective TF (ChIP-seq enrichment ranked by 20-20-20-15-10-10-5%, from weakest to strongest). Shaded areas represent standard deviation, and line colors represent fraction of significant response in the top bin (FDR < 0.05 & |log2 fold change| > 0.5). **d,** Representative locus showing that NRF1 binding upon DNA demethylation induces a distal TSS (500 nM DNMT1 inhibitor GSK-3484862 for 72 hours). **e,** Strength of distal TSS activation correlates with the NRF1 binding increase upon DNA methylation removal. NET-CAGE log2 fold change for TSS regions grouped by NRF1 binding changes upon DNA demethylation; bin labels indicate NRF1 binding fold change (log2, DNMT-TKO/WT) of the bin. **f,** Schematic and microscopy imaging of the mScarlet3 reporter. mScarlet3 signal is displayed with a 16-color lookup table. Cell colonies are outlined with gray dashed lines based on their nuclei staining. **g,** mScarlet3 transcription depends on the GABPA-driving TSS. To remove pre-existing proteins, mScarlet3 was bleached one hour following dTAG addition, and the newly produced mScarlet3 was quantified 12 hours after inducing degradation.

We tested whether de novo binding of a chromatin-opening TF is sufficient to trigger transcription initiation. To do so, we leveraged the methylation sensitivity of NRF1, which binds additional, primarily distal sites upon removal of DNA methylation in mESCs ^32^. NET-CAGE after global DNA demethylation using a DNMT1 inhibitor ^75^ revealed increased TSS signal predominantly at distal sites, most of which lacked pre-existing enhancer activity in untreated mESCs (**Extended Data Fig. 6f**). At such distal regions lacking prior enhancer activity (n = 2,201), a sharp increase in TSS signal coincided with increased NRF1 occupancy, exemplified by a representative distal NRF1 site (**Fig. 5d**). Genome-wide, the increase in TSS signal scaled with the gain in NRF1 binding following demethylation (**Fig. 5e**), suggesting that binding of a chromatin-opening TF can be sufficient to drive transcription initiation at distal sites.

We next examined whether TF-dependent distal initiation can function as a bona fide promoter and activate a proximal gene. We endogenously inserted a promoterless mScarlet3 coding sequence directly adjacent to a distal GABPA-bound site (**Fig. 5b,f**). This insertion induced robust expression of functional mScarlet3 protein, as detected by fluorescence microscopy (**Fig. 5f**). Targeted degradation showed that this promoter activity was highly dependent on GABPA at both protein (**Fig. 5g, Extended Data Fig. 6g,h**) and RNA levels (**Extended Data Fig. 6i**). Taken together, these results show that binding of chromatin-opening TFs at distal regions can be sufficient for transcription initiation of non-coding RNAs. Moreover, such TF-dependent distal initiation can confer functional promoter activity if a coding sequence is placed nearby.

## Discussion

Our comprehensive study uncovers a regulatory logic for CGIs, in which chromatin-opening TFs are the primary determinants of start site selection and thereby govern promoter output. Using a deep learning approach, we identified a core set of TFs (**Fig. 1**), many of which were previously implicated in promoter activation or transcription initiation ^45,76–80^. However, their roles had not been defined in a systematic, mechanistic framework. Reporter-based assays and in silico analyses ^77,78^ have suggested positional constraints of TF motifs, consistent with a set of TFs activating transcription downstream of their binding sites. Although these studies have provided important insights into promoter configuration, a causal and generalizable activation mechanism in genomic contexts remained unresolved. To address this gap, we combined endogenous TF perturbation with single-molecule footprinting and revealed the mechanism of promoter activation at the chromatin level. Our results revise the view that CGIs are inherently accessible and permissive ^6,19^ and instead show that CGI transcription is tightly regulated by a defined set of TFs that continuously open chromatin (**Fig. 2**).

At individual promoters, transcription is dominated by a single TF positioned closest to the dominant TSS, where it evicts nucleosomes and specifies the initiation site. Thus, TF contributions to gene expression are strongly shaped by motif position relative to other TFs. This argues against a purely additive model and instead supports a hierarchical architecture in which specific TFs dominate initiation at individual promoters ^81^. Removal of the dominant TF induces a TSS shift to an adjacent chromatin-opening TF (**Fig. 3**). This appears to be a common phenomenon at CGIs, as we observed TSS shifts at many promoters across TF perturbations. Given that relocation of the initiation site coincides with attenuated transcriptional responses to TF loss, we postulate that this mechanism contributes to stabilizing expression of broadly active essential genes against motif mutations or fluctuations in TF abundance.

Using integrated promoter variants, we establish that this TSS control is promoter-autonomous, therefore programmable in *cis*: manipulating chromatin-opening TF motifs repositions nucleosomes and relocates TSSs (**Fig. 4**). Together, our perturbations in *cis* and *trans* support a modular control of transcription initiation by chromatin-opening TFs as a common feature of CGI regulation. These findings establish a general function for chromatin-opening TFs in transcription initiation, extending beyond highly specialized factors such as NFYA ^76,82^.

Our findings do not exclude higher-order TF dependencies, which will require direct experimental testing such as combinatorial degradation of multiple TFs. Consistent with such interplay, at the *Rps21* promoter co-bound by GABPA and MYC, GABPA ablation nearly diminished transcription and accessibility, whereas MYC ablation only partially reduced transcription (**Fig. 4**). These results suggest that MYC-mediated activation depends on the presence of chromatin-opening TFs. Notably, MYC does not increase accessibility but instead amplifies transcription at promoters already opened by chromatin-opening TFs (**Fig. 2**). This scenario is consistent with reports that MYC promotes transcription elongation ^60,61^.

While the identified principle accounts for broadly distributed initiation at CGIs, we further show that chromatin-opening TFs similarly license TATA-box usage (**Fig. 4**). This argues that the TATA box functions as an auxiliary element that reinforces robust and precise TSS usage. Thus, chromatin-opening TFs can be sufficient to define transcription initiation, pointing to a minimalist architecture of mammalian promoters. These observations raise the possibility that transcription initiation, including pre-initiation complex loading, primarily requires open chromatin and presumably cofactors recruited by activating TFs. TF-mediated chromatin opening may also guide the use of other core promoter elements such as initiator (Inr) and downstream promoter element (DPE), which facilitate TFIID engagement ^26^.

Importantly, this principle of initiation control is not limited to promoters but also applies to enhancers, where it accounts for eRNA production (**Fig. 5**). By placing a coding sequence adjacent to an enhancer-associated TSS, we further show that binding of chromatin-opening TFs confers functional promoter activity, confirming a simple promoter architecture.

Together, our findings establish chromatin-opening TFs as a primary driver of transcription initiation. These results further suggest that mammalian promoters can operate with minimal sequence requirements – binding sites for chromatin-opening TFs. This sequence flexibility may help CGI promoters to retain robust activity despite their high mutation rate and rapid sequence evolution ^22^. The model further predicts that new initiation sites can emerge from relatively small sequence changes, potentially underlying rapid evolution of cis-regulatory elements, as observed for enhancers ^83^.

In conclusion, our findings reveal a chromatin-based mechanism of transcriptional activation across promoters and enhancers. This modular sequence grammar may support regulatory plasticity in gene expression and the rapid evolvability of *cis*-regulatory landscapes in mammalian genomes.

## Methods

### Cell culture

Mouse embryonic stem cells (mESCs, HA36 cell line) were cultured as previously described ^3^. Briefly, cells were maintained in ESC medium; Dulbecco’s modified Eagle’s medium (DMEM) (Thermo Fisher Scientific), supplemented with 15% fetal bovine serum (Thermo Fisher Scientific), 1× GlutaMax (Thermo Fisher Scientific), 1× nonessential amino acids (Gibco), 0.001% beta-mercaptoethanol (Sigma) and leukemia inhibitory factor (produced in-house). All experiments were performed with cells seeded on 0.2% gelatin-coated plates.

### Cell line generation

We endogenously tagged the target proteins with the degron tag peptide FKBP12(F36V), which allows inducible targeted protein degradation (dTAG system) ^53^. Gene fragments were synthesized (Twist Biosciences) containing the FKBP12(F36V) sequence and an epitope tag sequence (V5 or 3× FLAG tag) followed by a 2A peptide sequence and a selection marker sequence (blasticidin resistance gene, mNeonGreen or mScarlet3) flanked by homology arms around the insertion site of each target gene. Each end of the fragment contained a restriction enzyme cut site (ClaI or MluI) for cloning into a donor vector (a repair template vector). The fragment was also designed to contain a guide RNA (gRNA) sequence from the Escherichia coli genome (GTGTTGTGGACTGCGGCGGTCGG) for linearization of the repair template vector following nucleofection ^45^.

For the CRISPR-Cas9 protocol, we employed the Alt-R® CRISPR-Cas9 system (IDT). Briefly, to prepare 100 µM gRNA (crRNA:tracrRNA duplex), 200 µM crRNA (Alt-R® CRISPR-Cas9, IDT) targeting either the insertion site or the bacterial sequence was separately annealed with 200 µM tracrRNA (Alt-R® CRISPR-Cas9, IDT) by mixing them in a 1:1 ratio, heating at 95°C for 5 min, and cooling down to room temperature. Each annealed gRNA was mixed with an equal volume of Cas9 (Alt-R® S.p. Cas9 Nuclease V3, IDT) and incubated for 15 min at room temperature to form the ribonucleoprotein (RNP) complex. Then, 2 µL of each RNP complex (insertion site and bacterial sequence; total 4 µL) was mixed with 500 ng of the repair template vector, 0.4 µL of 100 µM electroporation enhancer (Alt-R® CRISPR-Cas9, IDT) and 2×10^5^ cells resuspended in nucleofection buffer (P3 Primary Cell 4D Nucleofector, Lonza), resulting in a 20 µL mixture. This mixture was electroporated using program CA120 of the Lonza 4D nucleofector. Immediately after electroporation, cells were transferred into culture medium supplemented with 0.1% homologous recombination enhancer (Alt-R® HDR Enhancer V2, IDT). The next day, the medium was exchanged with normal ESC medium, and cells were cultured for an additional two days.

For blasticidin selection, cells were treated with 8 µg/mL of blasticidin (InvivoGen) for 5 days, then seeded for clone picking and subsequent expansion. For the mNeonGreen or mScarlet3 selection, single cells were sorted by FACS based on the desired fluorescence signal. Successful tagging was verified by PCR and degradation of the target proteins was confirmed by western blotting (see the details in the section “Western blotting”).

We fused the degron tag to the C-terminus of all target TFs except the N-terminus of ATF1, because *Atf1* expresses multiple C-terminus isoforms. Notably, degron tagging for SP1 and ATF1 resulted in the successful tagging of only one allele and in the deletion of the other. The deletion in the *Atf1* allele is at the N-terminus, generating a null allele. The deletion in the *Sp1* allele is at the C-terminus and overlaps its zinc finger DNA binding domain. In line with this, expression of any truncated proteins corresponding to the deleted allele was not measured by western blotting using an antibody against the undeleted part of the protein, suggesting that the deleted allele is nonfunctional in both cases. Furthermore, complete degradation of these TFs under dTAG treatment was confirmed.

Regarding NRF1, we initially aimed to tag the N-terminus, because *Nrf1* expresses multiple C-terminus isoforms. However, cells with the N-terminus tagging led to increased expression of an isoform that skipped the degron-tagged N-terminus, leading to incomplete degradation of NRF1 protein. To enable the complete degradation of NRF1, alternatively two NRF1 isoforms whose C-terminus was fused to the degron tag were expressed from an ectopic locus using recombinase-mediated cassette exchange (RMCE) and the endogenous NRF1 was knocked out by deleting its fifth exon that overlaps its DNA binding domain using the Alt-R® CRISPR-Cas9 system. The RMCE insertion was performed as previously described ^64,65^. 4×10^6^ cells were transfected with 15 µg of pIC-Cre and 25 µg of L1-insert-1L vectors using Amaxa Nucleofection (Lonza). For the first isoform insertion, we selected cells with an insertion using ganciclovir (3 µM) starting two days after transfection for a total of 10 days. For the second isoform insertion, we used blasticidin (8 µg/mL) to positively select edited clones starting 3 days after transfection for a total of 5 days.

The entire panel of the degron cell lines was generated in this study except for the BANP degron cells, which were obtained from our previous study ^45^.

Regarding the mScarlet3 reporter knock-in for the GABPA binding distal TSS region, the same genome editing strategy as for the degron tag insertion was used. The cells expressing mScarlet3 were sorted by FACS and the genomic insertion was verified by PCR. The pool of cells with heterozygous or homozygous insertions was used for the experiments.

The DNA sequences used for generating the cell lines are listed in Supplementary Table 1.

### Western blotting

For nuclear cell lysis, 5×10^6^ cells per condition were first resuspended in 1 mL of lysis buffer (10 mM Tris-HCl pH 7.4, 10 mM NaCl, 3 mM MgCl2, 0.1 mM EDTA, 0.5% NP-40) supplemented with protease inhibitors (1× cOmplete Protease Inhibitor Cocktail, Roche) and incubated on ice for 10 min, followed by centrifugation at 3,000 rpm at 4°C for 5 min. The pellets were resuspended in 250 µL of wash buffer (10 mM Tris-HCl pH 7.4, 10 mM NaCl, 3 mM MgCl2, 0.1 mM EDTA) supplemented with protease inhibitors, and then centrifuged at 3,000 rpm at 4°C for 5 min. The resulting nuclei pellets were resuspended in 250 µL of TNN-lysis buffer (50 mM Tris-HCl pH 7.5, 250 mM NaCl, 0.5% NP-40, 5 mM EDTA, 1 mM DTT) with protease inhibitors, mixed briefly by vortexing and incubated on ice for 30 min. The samples were sonicated (2× 7 cycles of 30 sec ON / 30 sec OFF) using a Bioruptor Pico sonicator (Diagenode). Between each set of 7 cycles, the samples were incubated with 0.8 µL of benzonase (Sigma) at 12°C for 10 min. For GABPA and THAP11 degron cells, RIPA-lysis buffer (50 mM Tris-HCl, pH 8.0, 150 mM NaCl, 1% NP-40, 1% sodium deoxycholate, 0.1% SDS) was used instead of TNN-lysis buffer. For the NRF1 degron cells, whole cell lysis was performed using TNN-lysis buffer, followed by sonication and benzonase treatment. Finally, lysates were centrifuged at 13,000 rpm at 4°C for 15 min and the supernatant was collected for SDS-PAGE and western blotting.

Degradation of TFs was validated using antibodies raised against the endogenous peptides of target TFs as well as the epitope tag that was inserted alongside the degron tag. For ZFX, we did not find a suitable antibody, so its degradation was validated only by the antibody against the epitope tag. Of note, we observed instances of a small fraction of tagged protein that remained fused to the selection marker protein, suggesting that 2A-mediated cleavage is not fully efficient as reported ^84^. However, these proteins were fully degraded upon dTAG treatment. The antibodies used for Western blotting are listed in Supplementary Table 2.

### ChIP-seq

ChIP was carried out as previously described ^4^. 1.5×10^7^ mESCs were seeded onto 15 cm plates a day prior to the experiment. Cells were trypsinised, and after quenching the trypsinization with cell culture medium, formaldehyde was added for a final concentration of 1%. Cross-linking was performed for 10 min at room temperature.

From this point onwards, all steps were performed on ice, and centrifugations were done at 4°C. The reaction was quenched by adding glycine (125 mM final) for 10 min, followed by centrifugation at 600× g for 5 min. Cell pellets were rinsed once with cold PBS supplemented with protease inhibitors (1× cOmplete Protease Inhibitor Cocktail, Roche), followed by centrifugation at 600× g for 5 min. Pellets were resuspended first in 14 mL of buffer 1 (10 mM Tris pH 8.0, 10 mM EDTA, 0.5 mM EGTA and 0.25% Triton X-100), then in 14 mL of buffer 2 (10 mM Tris pH 8.0, 1 mM EDTA, 0.5 mM EGTA and 200 mM NaCl), with each resuspension followed by a 10-min incubation and centrifugation at 600× g for 5 min. Cells were then lysed in 1 mL HSB lysis buffer (50 mM HEPES pH 7.5, 500 mM NaCl, 1 mM EDTA, 1% Triton X-100, 0.1% sodium deoxycholate, 0.1% SDS and protease inhibitors) and incubated for 1 h. Lysates were sonicated using a Diagenode Bioruptor Pico (with 0.8 g sonication beads, 3× 10 cycles of 30 sec ON / 30 sec OFF) with a 5 min break on ice between each set of 10 cycles. Lysates were cleared by centrifugation at 12,000× g for 10 min before immunoprecipitation (IP). For the IP, lysates were first precleared with Protein A Dynabeads magnetic beads at 4°C for 1 h, followed by incubation with 5 µg antibody overnight at 4°C. The mixture was then incubated at 4°C for 3 h with Protein A Dynabeads magnetic beads. Beads were washed at room temperature twice with 1 mL HSB lysis buffer, once with 1 mL DOC buffer (10 mM Tris pH 8.0, 250 mM LiCl, 0.5% NP-40, 0.5% sodium deoxycholate and 1 mM EDTA). Beads were resuspended in 150 µL elution buffer (1% SDS, 0.1 M NaHCO3) followed by incubation with RNase A at 37°C for 30 min and with proteinase K at 65°C overnight. After reverse crosslinking, the eluted DNA was purified using Sera-Mag SpeedBeads (Cytiva).

IP was performed using the anti-FLAG tag antibody (Sigma, F1804) for ATF1 degron cells, the anti-NRF1 antibody (Abcam, ab175932) for NRF1 degron cells and the anti-V5 tag antibody (Thermo Fisher Scientific R960-25) for the rest of the cell lines. To control non-specific background signal, untagged WT mESCs were used for IP with anti-V5 or anti-FLAG antibodies for enrichment calculation (See the section “ChIP-seq analysis”). ChIP-seq for SP1 and c-MYC was performed using the single-TF-tagged cell lines.

The retrieved DNA fragments were prepared for sequencing using the NEBNext Ultra II DNA Library Prep Kit (NEB). The samples were amplified by PCR for 12 cycles. The libraries were sequenced using Illumina NovaSeq (2× 56bp paired-end). ChIP-seq experiments were performed in two independent replicates per condition.

### ATAC-seq

ATAC-seq was performed as previously described ^85,86^. mESCs were seeded onto six-well plates (2.0×10^5^ cells per well) a day prior to the experiment. Culture medium was replenished and supplemented with the dTAG compound (500 nM dTAG13, Sigma) or dimethyl sulfoxide (DMSO) for 4 h before harvest. For washing out experiments, after the first 4 h of treatment, cells were trypsinised and re-seeded on a new plate to culture medium without dTAG for another 16 h before harvest.

The cells were trypsinised and resuspended in cold PBS. 5×10^4^ cells were centrifuged at 500× g at 4°C for 5 min. Pellets were suspended in 50 μL RSB buffer (10 mM Tris-HCl pH 7.4, 10 mM NaCl and 3 mM MgCl_2_) containing 0.1% NP-40, 0.1% Tween-20 and 0.01% digitonin, then incubated on ice for 3 min. Afterwards, 1 mL of RSB buffer containing 0.1% Tween-20 (without NP-40 or digitonin) was added, followed by centrifugation at 600× g at 4°C for 10 min. After discarding the supernatant, nuclei were isolated and resuspended in 50 μL of Tn5 transposition mix: 10 μL of 5× reaction buffer (50 mM Tris-Cl pH 8.5, 25 mM MgCl_2_, 50% dimethylformamide) supplemented with 2.5 μL of transposase (100 nM final), 16.5 µL of PBS, 0.5 µL of 1% digitonin, 0.5 µL of 10% Tween-20 and 20 µL of water. The transposition reaction was performed in a thermomixer shaking at 300 rpm at 37°C for 30 min. The tagmented DNA was purified using MinElute PCR Purification Kit (Qiagen) and was eluted in 20 μL. The samples were amplified by PCR using Q5 High-Fidelity Polymerase (NEB) for 5 cycles. The libraries were sequenced using Illumina NovaSeq (2× 56bp paired-end). ATAC-seq experiments were performed in two independent replicates per condition.

### RNA-seq

2.0×10^5^ mESCs per well were seeded onto twelve-well plates a day prior to the experiment. Where applicable, culture medium was replenished with or without dTAG(500 nM dTAG13, Sigma) 8 h before harvesting with QIAzol. RNA was isolated using a Direct-zol MicroPrep RNA Purification Kit (Zymo), following the manufacturer’s instructions. Sequencing libraries were prepared using Illumina Stranded total RNA-seq Prep kit and were sequenced using Illumina NovaSeq (2× 56bp paired-end). RNA-seq experiments were performed in two independent replicates per condition.

### NET-CAGE

#### Nascent RNA collection

Nascent RNA extraction was performed as previously described ^62^. 5×10^6^ mESCs were seeded on 10 cm plates a day prior to the experiment. Where applicable, culture medium was replenished with or without dTAG (dTAG13 500 nM, Sigma) for 4 h. For DNA demethylation, the cells were treated with DNMT1 inhibitor (500 nM GSK-3484862, Merck) ^75,87^ or DMSO for 72 h.

The cells were trypsinised, collected, and washed once with PBS. After the wash, the cell pellets were resuspended in 1 mL of Buffer A (Nuclei EZ Lysis Buffer (Sigma) supplemented with 25 μM α-amanitin (Sigma), 1× cOmplete Protease Inhibitor Cocktail (Roche) and SUPERase-IN RNase Inhibitor (20 U, Thermo Fisher Scientific)), incubated on ice for 10 min and centrifuged at 800× g at 4°C for 5 min. After discarding the supernatant, the pellets were washed again with 600 µL of Buffer A. The washed pellets were resuspended in 200 µL of Buffer B (1% NP-40 (Thermo Fisher Scientific), 20 mM HEPES pH 7.5, 300 mM NaCl, 2 M urea, 0.2 mM EDTA, 1 mM dithiothreitol (DTT), 25 μM α-amanitin, 1× cOmplete Protease Inhibitor Cocktail (Roche) and SUPERase-IN RNase Inhibitor (20 U, Thermo Fisher Scientific)), incubated on ice and centrifuged at 3000× g at 4°C for 4 min. After discarding the soluble fraction, the remaining insoluble fraction was washed again with 100 µL Buffer B. The resulting fractionated chromatin was treated with 50 µL DNase I solution (10U, Thermo Fisher Scientific) for 30 mins at 37°C. The final material was directly added to 700 µL QIAzol, RNA was extracted using miRNeasy Micro kit (QIAGEN), following the manufacturer’s instructions. RNA was eluted in 12 µL nuclease free water (NFW) and quantified by Qubit RNA Broad Range Kit (Thermo Fisher Scientific). NET-CAGE experiments were performed in two independent replicates per condition.

#### CAGE library preparation

2.5 µg of RNA was used for CAGE library preparation as previously described ^88,89^ with the following modifications. After the biotinylation of the 5’ cap and pulldown, the cDNA was ligated to linkers that were compatible with NEBNext Multiplex Oligos for Illumina. After linker ligation to both ends, the second strand of the cDNA was performed in a 50 µL reaction (40 µL cDNA solution, 2 µL NFW, 1 µL of 10 mM dNTPs, 1 µL Deep Vent (exo-) DNA polymerase (2 U/µL, NEB), 5 µL ThermoPol Reaction Buffer pack and 1 µL of 50 µM second strand primer). The mixture was incubated with the following program: 95°C for 5 min, 55°C for 5 min and 72°C for 30 sec and 4°C to hold. The samples were amplified in a 50 µL reaction of 15 µL cDNA solution, 25 µL Q5 High-Fidelity Polymerase (NEB) and the 10 µL Index primer mix with the following PCR program: 98°C for 30 sec, 9 cycles of (98°C for 10 sec and 65°C for 75 sec), 65°C for 5 min and 4°C to hold. The libraries were sequenced using Illumina NovaSeq (2× 56bp paired-end). The DNA sequences used for CAGE library preparation are listed in Supplementary Table 3.

### Locus-specific CAGE

#### Library preparation

RMCE insertion was performed as previously described ^64^. 4×10^6^ mESCs were transfected with 15 µg of pIC-Cre and 25 µg of the pool of promoter inserts using Amaxa Nucleofection (Lonza). The constructs were designed to contain promoter variant sequence coupled to a luciferase coding sequence followed by DNA barcodes unique to each promoter. The DNA sequences of the promoter inserts are listed in Supplementary Table 4. The cells were treated with ganciclovir (3 µM) starting 2 days after transfection for a total of 10 days and then harvested for RNA extraction. The total RNA was extracted using miRNeasy Micro kit (QIAGEN), following the manufacturer’s instructions.

To specifically reverse-transcribe transcripts from the integrated promoters, 2.5 µg of RNA in 9 µL of NFW was mixed with 1 µL of a 2 µM reverse transcription (RT) primer encoding a unique molecular identifier (UMI). The RNA with the RT primer was denatured at 65°C for 5 min and was immediately cooled down on ice. The 10 µL of denatured RNA was mixed on ice with 10 µL of RT reaction buffer (400 U Super Script III Reverse Transcriptase (Thermo Fisher Scientific), 2× First-strand buffer, 10 mM DTT and 4 mM dNTPs (1 mM each)) into the 20 µL final volume. The mixture was incubated at 55°C for 60 min. The cDNA-RNA hybrid was purified with 1.8× RNAClean XP beads (Beckman Coulter) and was eluted in 40 µL of NFW. The 40 µL of cDNA-RNA hybrid was added to 4 µL diol oxidation buffer (2 µL of 1 M NaOAc pH 4.5 and 2 µL of 250 mM NaIO4) and incubated on ice protected from light for 5 min. Then, 16 µL of 1 M Tris-HCl (pH 8.5) was added to stop the reaction. The oxidized cDNA-RNA hybrid was purified with 1.8× RNAClean XP beads (Beckman Coulter) and was eluted in 40 µL of NFW. The 40 µL cDNA-RNA hybrid was added to 8 µL biotinylation buffer (4 µL of 1 M NaOAc pH 6.0 and 4 µL of 10 mM biotin hydrazide) and incubated at 23°C for 2 h. The biotinylated cDNA-RNA hybrid was purified with 1.8× RNAClean XP beads (Beckman Coulter) and was eluted in 40 µL of NFW. The 40 µL biotinylated cDNA-RNA hybrid was added to 5 µL RNase I reaction buffer (0.5 µL RNase ONE (10 U/µL, Promega) and 4.5 µL of 10× RNase ONE buffer) and was incubated at 37°C for 30 min. The reaction mixture was subsequently purified using 1.8× RNAClean XP beads (Beckman Coulter) and was eluted in 40 µL of NFW.

To prepare streptavidin beads for the biotinylated cDNA-RNA hybrid, 30 µL of Dynabeads M-270 Streptavidin beads (Thermo Fisher Scientific) per sample were added to a new 1.5 mL tube and set on a magnetic stand. The beads were washed twice with equal volumes of LiCl buffer (20 mM Tris-HCl pH 7.5, 7 M Lithium chloride, 0.1% Tween-20, 2 mM EDTA pH 8.0) by magnetizing the beads and discarding the supernatant.

Finally, 95 µL of LiCl buffer per sample were added to the beads and kept on ice until used. The 95 µL of the prepared beads was added to the 40 µL of cDNA-RNA hybrid and incubated for 15 min at 37°C. The beads were washed with 150 µL of TE wash buffer (10 mM Tris-HCl pH 7.5, 0.1% Tween-20, 1 mM EDTA pH 8.0) three times in total. After discarding the supernatant, 35 µL of release buffer (1× RNase ONE Reaction buffer (Promega) and 0.01% Tween-20) was added to the beads and the beads were incubated at 95°C for 5 min. Then, the samples were immediately cooled down on ice and set on the magnetic stand. The 35 µL of cDNA in the supernatant was collected into a new tube. To completely collect the cDNA, another 30 µL of release buffer was added to the beads and again the supernatant was collected. The resulting 65 µL of cDNA was added to 5 µL of RNase reaction buffer (2.9 µL of release buffer, 2 µL of RNase ONE ribonuclease (10 U/µL, Promega) and 0.1 µL of RNase H (60 U/µL, Takara)) and incubated at 37°C for 30 min. The mixture was then purified with 1.8× AMPure XP beads (Beckman Coulter) and eluted in 25 µL of NFW.

The cDNA was completely dried up using a centrifugal concentrator (SpeedVAC) and added to 4 µL NFW. The 4 µL of cDNA was incubated at 95°C for 5 min and immediately cooled down on ice. Separately, 4 µL of 5’ linker (*) (2.5 µM) was incubated at 55°C for 5 min and immediately cooled down on ice. The 4 µL of cDNA and the 4 µL of 5 linker were added to 16 µL of DNA ligation Mighty Mix (Takara) on ice. The cDNA was incubated at 16°C for 16 h. *\* 5’ linker was prepared by annealing 100 µM of each oligo in linker buffer (TE buffer with final 100 mM NaCl) and diluted 40× using the same linker buffer (2.5 µM final). The DNA sequences of oligos for 5’ linker are listed in Supplementary Table 5*.

The resulting ligated DNA was purified with 1.8× AMPure XP beads (Beckman Coulter) twice to completely remove the remaining linker and was eluted in 40 µL of NFW. The cDNA was added to 10 µL of phosphatase reaction buffer (1 µL of Shrimp Alkaline Phosphatase (1 U/µL, Thermo Fisher Scientific), 5 µL of 10× SAP buffer and 4 µL of NFW) and was incubated at 37°C for 30 min followed by 65°C for 15 min and 4°C to hold.

Then, 2 µL of USER enzyme was added to the SAP treated cDNA and the mixture was incubated at 37°C for 30 min, followed by 95°C for 5 min. After incubation, the tube was immediately cooled down on ice, purified with 1.8× AMPure XP beads (Beckman Coulter) and eluted in 40 µL of NFW. The cDNA was added to 10 µL of second strand synthesis buffer (2 µL of NFW, 1 µL of 10 mM dNTPs, 1 µL Deep Vent (exo-) DNA polymerase (2 U/µL, NEB), 5 µL ThermoPol Reaction Buffer pack and 1 µL of 50 µM second strand primer) and the second strand was synthesized with the following program: 95°C for 5 min, 6 cycles of (95°C for 30 sec, 55°C for 30 sec and 72°C for 1 min), 72°C for 5 min and 4°C to hold. Then, 1 µL of Exonuclease I (NEB) was added to the samples and was incubated at 37°C for 30 min to remove the remaining primer. The samples were purified with 1.8× AMPure XP beads (Beckman Coulter) twice to completely remove Exonuclease I and were eluted in 15 µL of NFW. The cDNA was mixed with 25 µL Q5 High-Fidelity Polymerase (NEB) and 10 µL of the Index primer mix (NEBNext Multiplex Oligos for Illumina) and amplified with the following PCR program: 98°C for 30 sec, 20 cycles of (98°C for 10 sec and 65°C for 75 sec), 65°C for 5 min and 4 °C to hold. The resulting 60 µL of the library was purified with 1.2× AMPure XP beads (Beckman Coulter) and eluted in 12 µL of NFW. The libraries were sequenced using Illumina NovaSeq (2× 56bp paired-end). The DNA sequences used for locus-specific CAGE are listed in Supplementary Table 4,5. Locus-specific CAGE experiments were performed in two independent replicates.

### Barcoded DNA sequencing for estimation of insertion ratio

To estimate the ratio of inserts in the pool of the cells used for locus-specific CAGE, sequencing for barcoded DNA amplicons was performed. After ganciclovir selection, genomic DNA (gDNA) was extracted from 5×10^6^ cells using the Quick-DNA miniprep plus kit (Zymo) following the manufacturer’s instructions. The extracted gDNA was amplified using primers specific to the RMCE inserts in 50 µL of PCR reaction of 1 µg gDNA, 25 µL of Q5 High-Fidelity Polymerase (NEB), 1 µM of each primer with the following PCR program: 98°C for 3 min, 12 cycles of (98°C for 10 sec and 65°C for 75 sec), 65°C for 5 min and 4°C to hold. The amplified barcoded DNA was purified using AMPure XP beads (Beckman Coulter) and was eluted in 15 µL of NFW. The primers for the initial PCR contain overhanging sequences that were used as the primer binding sites for the second PCR amplification. This second PCR was performed in the reaction of 15 µL of barcoded DNA, 25 µL Q5 High-Fidelity Polymerase (NEB), the 10 µL Index primer mix and amplified with the following PCR program: 98°C for 30 sec, 18 cycles of (98°C for 10 sec and 65°C for 75 sec), 65°C for 5 min and 4°C to hold. The libraries were sequenced using Illumina NovaSeq (2× 56bp paired-end). The primers used to amplify the barcoded DNA are listed in Supplementary Table 5.

### Single-molecule footprinting

Single-molecule footprinting was performed as previously described ^65,90^ with several modifications. After ganciclovir selection, 2.5×10^5^ cells were collected, washed with cold PBS and incubated for 10 min on ice in 1 mL of ice-cold lysis buffer (10 mM Tris pH 7.5, 10 mM NaCl, 3 mM MgCl_2_, 0.1 mM EDTA and 0.5% NP-40) to extract their nuclei.

After centrifugation at 800× g at 4°C for 5 min, the nuclei were washed once with 250 µL of ice-cold wash buffer (10 mM Tris pH 7.5, 10 mM NaCl, 3 mM MgCl_2_ and 0.1 mM EDTA) and followed by centrifuged again at 800× g at 4°C for 5min. The samples were then resuspended in 100 µL of 1× M.CviPI buffer, mixed with 200 µL of GpC methyltransferase reaction mix (200 U M.CviPI (NEB), 1× M.CviPI buffer, 1 mM SAM, 300 mM sucrose) on ice and then incubated at 37°C for 15 min. The reaction was stopped by adding 300 µL of Stop Solution (20 mM Tris-HCl pH 8.0, 600 mM NaCl, 1% SDS, 10 mM EDTA) pre-warmed at 37°C. To remove proteins and RNA, the samples were treated with 12 µL of proteinase K (10 mg/mL) at 65°C for 16 h and then with 10 µL of RNase A (10 mg/mL) at 37°C for 30 min. Finally, gDNA was extracted with phenol-chloroform purification and isopropanol precipitation as follows. The samples were mixed with 600 µL of a 1:1 phenol-chloroform solution, vortexed at maximum speed for 1 min and centrifuged at 10,000× g for 10 min. The supernatant was then transferred to new tubes and mixed with 600 µL of chloroform, vortexed and centrifuged again as before. The supernatant was transferred to new tubes and mixed with 600 µL of isopropanol and 1 µL of glycogen and incubated at RT for 10 min followed by centrifugation at maximum speed at 4°C for 1 h. The pellet was then washed with 500 µL of ice-cold 70% ethanol and centrifuged at maximum speed at 4°C for 15 min. The supernatant was discarded, and the pellet was dried at 37°C for 15 min and resuspended in 22 µL NFW at RT overnight. After quantification, 2 μg of the gDNA was converted using the EZ DNA methylation-gold kit (Zymo) following the manufacturer’s instructions. Target regions were amplified from bisulfite-converted DNA using KAPA HiFi HotStart Uracil+ReadyMix Kit (Roche) with the following program: 95°C for 4 min, 35 cycles of (98°C for 20 sec, 60°C for 15 sec and 72°C for 30 sec), 75°C for 5 min and 4°C to hold. The amplicons were then purified with 0.6× AMPure XP Beads and used as input for library preparation. The retrieved amplicons were prepared for sequencing using the NEBNext Ultra II DNA Library Prep Kit (NEB). The samples were amplified by PCR for 7 cycles. The libraries were sequenced using Illumina MiSeq or MiSeq i100 (600bp paired-end). The DNA sequences used for bisulfite-converted DNA amplification as well as the promoter sequences are listed in Supplementary Table 4,6. Single-molecule footprinting experiments were performed in two to four independent replicates.

### Quantification of distal TSS reporter by light microscopy imaging and RT-qPCR

mESCs expressing an mScarlet3 reporter at the GABPA-bound distal TSS were obtained as described in the section “Cell line generation” and were cultured on gelatin-coated 8-well μ-slides (Ibidi). For GABPA degradation, cells were treated with either DMSO or dTAG13 (500 nM dTAG13, Sigma) for 12 h. Fluorescence recovery after photobleaching was performed using a Yokogawa CSU-W1 spinning disk confocal microscope equipped with a photokinesis unit. One hour after initiating treatment (DMSO or dTAG), cells were photobleached using a 488 nm laser with a 40× objective. Following 12 h of treatment, nuclei were stained with Hoechst, and cells were fixed with 4% paraformaldehyde (PFA). mScarlet3 and Hoechst fluorescence images were acquired using the spinning disk confocal microscope with a 40× objective. Images were processed using ImageJ. Each treatment condition was analyzed in duplicate. Nuclear mean fluorescence intensity of mScarlet3 was quantified for individual cells.

Regarding the RT-qPCR, the cells were treated with either DMSO or dTAG (500 nM dTAG13, Sigma) for 8 h before RNA collection in the same way as for RNA-seq experiments. The RT-qPCR was performed using the PrimeScript RT kit (Takara) and the DNA sequences for the primers are listed in Supplementary Table 7.

### Computational analysis

#### Annotations and datasets for epigenetic signatures

Genome-wide annotations for promoters and CpG islands (CGIs) were obtained from UCSC as previously described ^65^. Briefly, promoters were retrieved from the TxDb.Mmusculus.UCSC.mm10.knownGene (v3.10.0) package, and CGI annotations were imported from the ‘cpgIslandExt’ table using rtracklayer (v 1.68.0) ^91^. All regions were filtered against the mouse ENCODE blacklisted regions ^92^. For UCSC genes annotated with multiple TSSs, we retained the one with the highest Pol II ChIP-seq enrichment signal relative to input as measured from GSM747548 and GSM747546 ^93^ (see ChIP-seq analysis). Promoters extending from −800 to +200 bp relative to the TSS were classified as CGI promoters if they overlapped a CGI. UCSC TSS annotations were used as the initial reference for promoter-associated analyses and were later refined by high-resolution TSS definitions derived from our NET-CAGE profiles in mESCs. For genomic intervals including promoters, DNase I hypersensitivity (DHS) was quantified using the qCount (log2 with a pseudocount of 8) function available from QuasR (v1.48.1) ^94^ package utilizing the publicly available DNase-seq dataset for mESCs (GSE67867 ^32^). We compared accessibility only among active CGI promoters, defined as those with RPKM ≥ 1 in WT mESCs (see “RNA-seq analysis” for gene expression quantification). To minimize potential confounding effects of DNA methylation in inactive CGIs, accessibility analysis was further restricted to CGI promoters with < 20% DNA methylation levels quantified from GSM6283069 ^43^.

### Convolutional neural network analysis

We implemented a sequence-to-activity model using a convolutional neural network (CNN) following the framework previously described in Durdu et al. ^37^. In brief, the CNN was trained on DNA sequences derived from CGI regions to predict the continuous values of DNase-seq enrichment, which served as a proxy for chromatin accessibility. For model input, CGI intervals were partitioned into 150-bp non-overlapping windows and CGIs overlapping with the CGI promoters were position-shifted relative to the TSS. Input DNA sequences were encoded in a one-hot representation over 150 bp windows. The CNN model was trained and evaluated using both the positive and negative strands of each genomic locus. To generate the model’s training targets, log2-transformed DHS values were quantified as described above. DHS values were truncated at a lower bound of 6 and subsequently shifted to remove the pseudocount baseline. The CNN design was based on the Basset framework ^36^ and further adapted using the architectural choices and hyperparameters of DeepSTARR ^95^. The final architecture comprised four consecutive 1D convolutional layers with 512, 512, 512 and 256 filters and kernel sizes of 12, 3, 5 and 3, respectively. Each convolutional layer was followed by a ReLU activation and max-pooling with a pool size of 2. These layers fed into two fully connected layers with 512 and 256 units, both using ReLU activation and a dropout rate of 0.4. The output layer employed a linear activation function to predict the DHS accessibility values in mESCs. Model development was carried out in Keras ^96^ via the KerasR package (v2.2.5.0) ^97^ using a TensorFlow backend (v2.0.0) ^98^. We optimized the network with the Adam optimizer and a mean squared error (MSE) loss function, using a batch size of 64. Training employed early stopping based on a 20% validation split from the training set, with a patience of 15 epochs. 150-bp non-overlapping CGI-windows on chromosome 19 were held out entirely for testing and excluded from both training and validation. In addition to MSE-based evaluation, we assessed model performance on the held-out test chromosome (chromosome 19) using Pearson’s correlation. 150 bp CGI-windows on chromosome 18 were used as a validation set for hyperparameter tuning.

To interpret the CNN model, we applied the shap.DeepExplainer function from the SHAP library ^99^, which provides an adapted implementation of the DeepLIFT algorithm^38^. This approach provided nucleotide-level contribution scores for all 150-bp non-overlapping CGI-windows. As background input for DeepExplainer, we generated 100 dinucleotide-shuffled versions for each CGI-window. For clustering and summarizing sequence patterns discovered by the CNN, we input the hypothetical/contribution scores into TF-MoDISco Lite ^39^ (https://github.com/jmschrei/tfmodisco-lite), using a sliding window length of 15 bp, additional flanks of 5 bp, and a target seqlet FDR cutoff of 0.15. We retained only TF-MoDISco metaclusters with more than 300 seqlets that also showed significant similarity to a HOCOMOCO motif (*q-value* < 0.01). For contribution weight matrices, the hypothetical/contribution scores corresponding to the one-hot encoded DNA sequences were used and visualized with ggseqlogo (v0.2)^100^. For each CGI window, all occurrences of the TF motifs were mutated, and the resulting sequences were evaluated using the DL model. Motif instances selected for mutagenesis were defined from an independent set of sites derived from ChIP-seq binding profiles (see ‘*de novo* motif search’).

### Motif enrichment analysis at CGIs

As an orthogonal analysis to the deep learning (DL) framework, motif enrichment was carried out using the JASPAR 2018 motif database ^101^ implemented in the R package (v1.1.1). To reduce redundancy among highly similar motifs, we used non-redundant motif collections to refine motif logos ^102^. BANP and THAP11/ZNF143 motifs, which are not included in these collections, were added manually ^45,103^. Motif enrichment among active CGI promoters (RPKM > 1) was performed with the monaLisa R package (v1.14.1) ^46^ using the mm10 reference genome as background and requiring a minimum motif match score of 10 for matches for the non-redundant motif library. To investigate sequence determinants underlying differences in CGI accessibility, we further performed motif enrichment comparing highly accessible versus lowly accessible CGIs at active promoters (DHS > 9 vs DHS < 9). Differential motif enrichment between these two bins was carried out with monaLisa using a minimum motif match score of 10 (argument “min.score”) in the calcBinnedMotifEnrR function.

### ChIP-seq analysis

#### Mapping and counting reads

ChIP-seq datasets were aligned to the mm10 assembly of the mouse genome using the Bioconductor package QuasR ^94^, which internally uses Bowtie (via the RBowtie package) ^104^. Bowtie was run with QuasR default parameters, allowing only for uniquely mapped reads.

Read counts over defined genomic regions were generated using the QuasR function qCount, with the selected options (shift = “halfInsert” and useRead = “first”). The counts were normalized between the datasets by library size, scaling the total read counts to that of the minimum library size. The normalized counts were then log_2_ transformed after adding a pseudocount of 8. Enrichment of log_2_ ChIP-seq read counts was calculated by subtracting the matched log_2_ counts of the corresponding control datasets, which were the ChIP-seq samples using the respective epitope tag antibody (V5/Flag) against untagged mESCs. ChIP-seq datasets for NRF1 in a DNMT-TKO background, BANP and NFYA were collected from published works ^32,45,105^.

#### Binding analysis for peaks and de novo motif search

Peak calling was performed with MACS2 (v.2.2.7) ^106^ using the callpeak function with default settings and specifying the genome size with-g mm. Peaks were called for each replicate, separately. Resulting peaks were then filtered for those that overlapped at least 70% between replicates and filtered against the ENCODE blacklisted regions ^92^.

The resulting reproducible peaks were then used to count the reads over a ±100 nt region centered on the peak summit position (midpoint of peak summit points from two replicates) to calculate the enrichment as described above. For each TF, peaks were sorted by log_2_ enrichment, and the top 1,000 peaks were used for de novo motif finding using HOMER ^107^. HOMER was run using the function findMotifsGenome.pl for 10 bp length (-len 10). For the TFs that were known to have longer motifs, HOMER was run for 12 bp (BANP, NRF1 and SP1) or 20 bp (THAP11). For each dataset, the top motif was retrieved and flanking bases with low information content were removed and visualized using the Bioconductor R package ggseqlogo ^100^. Based on these de novo motifs, we predicted genomic locations of the motifs as previously described ^45^, using a log-odds score cut-off (position weight matrix score > 9). To define threshold values for ChIP-seq enrichment at peaks to select binding sites, we calculated the frequency of motif occurrence per peak as a function of binding enrichment. More precisely, we calculated the frequency of motif occurrence per peak by calling all predicted motif sites that overlap each peak (±100 nt region centered on peak summit position) and defined the enrichment value threshold for “binding” by the minimum value where the majority (>50%) of the peaks contain at least one respective motif (indicated in **Extended Data Fig. 1f**). This allows us to control differences in dynamic ranges of binding enrichment of each ChIP-seq experiment. The resulting peaks above the enrichment threshold were selected as binding peaks.

#### Binding strength analysis

ChIP-seq reads were counted over a 1,001 nt window (-800/+200) relative to TSSs (defined by either UCSC annotation in **Fig. 1 and 2** or our NET-CAGE datasets in **Fig. 3 to 5**, see the section “NET-CAGE defined TSS” for the promoter definition) and the resulting binding enrichment at each region was used to show the continuous scale of binding strength. To define bound TSS regions for each TF, we selected the TSS regions (1,001 nt) that overlapped with a summit position of binding peaks as defined above.

#### Binding analysis for motifs and their position

To analyze TF binding on motifs, we used the genome-wide predicted motif location as described above. To define bound motifs by each TF, we selected those predicted motifs that overlapped with ±100 nt windows centered on peak summit positions of binding peaks as defined above. These bound motifs were used to identify the TF binding positions at a given promoter. To define TSS-proximal TFs, the nearest bound motif of each TF was identified within-800 to 0 bp upstream of TSSs defined by NET-CAGE. The nearest bound motif among chromatin-opening TFs was called as the most proximal TF.

### ATAC-seq analysis

ATAC-seq reads were trimmed using cutadapt (v.1.18) ^108^, with parameters-a CTGTCTCTTATACACA-A CTGTCTCTTATACACA-m 5-overlap = 1 and then mapped to mm10. The resulting datasets were aligned to mm10 using QuasR as for ChIP-seq.

Read counts over a 201 nt window centered on TF motifs or a 501 nt window (-400/+100) relative to TSSs were generated using the QuasR function qCount, with default parameters and without shifting (to quantify insertion sites). The resulting read counts were then normalized by edgeR (v.4.2.0) ^109^ calcNormFactors function with the default parameters and fold change estimates were derived using the edgeR predFC function with a pseudocount of 8. For each differential analysis comparison, regions were considered to be differentially accessible with a false discovery rate (FDR) < 0.05 & an absolute log2 fold change > 0.5.

Meta profiles and heatmap profiles for accessibility changes were generated using QuasR function qProfile and normalized by scaling the total mapped reads to 30e6 reads. Profiles were then smoothed with a running mean of 51 nt and the mean signal of replicates are shown in figures.

### RNA-seq analysis

RNA-seq reads were mapped to mm10 using QuasR in spliced alignment mode (splicedAlignment = TRUE) with the option (aligner = “Rhisat2”), which internally uses HISAT2 ^110^. Gene expression levels were calculated using the QuasR function qCount (with the options orientation = “opposite” and useRead = “first”) using the University of California Santa Cruz (UCSC) Known Genes database, which was accessed via the Bioconductor package TxDb.Mmusculus.UCSC.mm10.knownGene [https://bioconductor.org/packages/release/data/annotation/html/TxDb.Mmusculus.U <u>CSC.mm10.knownGene.html</u>] with the option reportLevel = “gene”, which counts the total number of reads mapping to any base annotated as exonic for each gene. The resulting read counts at the gene level were used to calculate RPKM and gene expression change. For the gene expression change, fold change estimates were derived using the edgeR predFC function with a pseudocount of 8. For each differential analysis comparison, genes were considered to be differentially expressed with a false discovery rate (FDR) < 0.05 & an absolute log2 fold change > 0.5 to define moderate to strong gene expression response.

For UCSC-defined promoters, active promoters were defined as having RPKM >= 1 in WT mESCs. For the heatmap in **Fig. 2c**, the log2 fold changes of gene expression between control and TF-degraded from all tested degron cells were clustered using *k*-means clustering with *k = 11.* The number of clusters, *k*, was determined by a value over which reduction in the total within-cluster sum of squares appeared less significant.

For each promoter, the maximum downregulation of gene expression among all TF degradation conditions was chosen and within each cluster, promoters were sorted by this value from more downregulation (left) to less downregulation (right). The fold changes for RNA-seq and ATAC-seq, and TF binding enrichment at promoters of ChIP-seq from each degron cell line datasets were visualized using ComplexHeatmap (v2.2.6) ^111^.

### NET-CAGE

#### NET-CAGE defined TSSs

To select the reads coming from 5’ capped RNA, a R package ShortRead (v.1.68.0) function readFastq ^112^ was used and the reads that start from G (methyl-guanosine of 5’ cap) were extracted. Using the resulting fastq files, reads were mapped using STAR (v 2.7.9) ^113^. The mapped reads were filtered to retain only read1 (representing 5’ of transcripts) and used to define the transcription start sites by CAGEr (v.2.14.0) ^114^. Here, CAGEr was run together for all the datasets collected in this study (with/without perturbation). TSS clusters were called by the CAGEr function distclu with options maxDist = 100 and keepSingletonsAbove = 5. Consensus clusters were called by aggregateTagClusters with the default parameters, resulting in n = 30,869 consensus TSS clusters. TSS width was calculated by quantilePositions with the default parameters. These NET-CAGE defined consensus TSS clusters were annotated for the nearest gene name and distance by the ChIPseeker (v1.34.1) ^115^ function annotatePeak with the options tssRegion = c(-800,200) using TxDb.Mmusculus.UCSC.mm10.knownGene. The TSS regions were set to the position of dominant TSSs from WT mESCs data. The TSS regions were filtered against ENCODE blacklist, resulting in 30,785 NET-CAGE TSS regions.

NET-CAGE–defined TSSs were further characterized using previously established chromatin-state maps for mESCs ^116^ (downloaded from https://github.com/guifengwei/ChromHMM_mESC_mm10). These genome-wide maps were generated by ChromHMM ^117,118^ using ENCODE ChIP-seq profiles of histone marks and core regulatory factors ^119^. For TSS regions overlapping with multiple chromatin states, assignment was prioritized in the following order: “Active Promoter:E7”, “Bivalent Promoter:E6”, “Transcriptional transition:E9”, “Strong Enhancer:E8”, “Enhancer:E4”, “Weak/Poised Enhancer:E11”, “Repressed Chromatin:E5”, “Heterochromatin:E3”, “Transcriptional elongation:E10”, “CTCF(Insulator):E1”, “Intergenic Region:E2”, “Low Signal/Repetitive Elements:E12”.

To define promoters, we selected TSS regions that are annotated as “promoter” by CAGEr. (n =18,086). To generate a non-redundant promoter set, gene annotation by ChIPseeker was used to find promoters that share the same gene (alternative promoters). In these cases, a promoter with the highest NET-CAGE signal (tpm) per gene was selected, enabling the identification of the dominant promoters as well as comparison of NET-CAGE data with RNA-seq data. To select the reliably expressed TSS in mESCs, we used mean tpm > 5 in WT mESCs, resulting in non-redundant, 12,631 active promoters in mESCs that were used for promoter analysis.

To define distal TSSs, we filtered out TSS regions that were annotated as “promoter” by CAGEr or those overlapping with NET-CAGE defined promoters (-800/+200 bp relative to TSSs) and only selected the ones that were located at least 1,000 bp distant from the TSS of mm10 genes using the calculation (distanceToTSS) by ChIPseeker, resulting in a set of promoter-distal TSS regions (n = 11,137). Following this, the regions that showed enhancer chromatin signatures were selected. Here, the regions that overlapped enhancer-like regions (E4, E8) defined by the ChromHMM maps as well as H3K27ac peaks in mESCs from our previous study (GSM1891651) ^32^ were selected. To select the reliably expressed TSS in mESCs, we used mean tpm > 5 in WT mESCs, resulting in 2,640 candidate enhancers. Regarding TSS analysis upon DNMT1 inhibition, in order to filter out distal TSS regions with pre-existing cis-regulatory element (CRE) signature in WT mESCs, the promoter-distal TSS regions (n = 11,137) were selected for the ones that did not overlap regions with promoter or enhancer signatures (E4, E6, E7, E8, E9, E11) defined by the ChromHMM maps as well as H3K27ac or H3K4me1 peaks. This resulted in 2,201 TSS regions without pre-existing CRE signatures, which were used only for analysis of NRF1 effect upon DNA demethylation.

#### TATA-box promoters

The canonical TATA box motif was obtained from JASPAR2020 (TBP MA0108.2) ^120^. The NET-CAGE defined promoters that contained a TATA box at-35 to-21 bp upstream of the TSS were selected as TATA-box promoters.

#### TSS shift analysis

The TSS shifts were called using the CAGEr function scoreShift for all TSS regions. Here, CAGEr systematically detects spatial separation of TSS usage by comparing cumulative distribution of TSS signal along the same consensus TSS cluster region between two groups (control and dTAG treatment). Using the resulting scores, TSS shifts were called with the threshold shifting.score >= 0.2 and fdr.KS (adjusted *P*-value from Kolmogorov-Smirnov test) <= 0.01. This function also provides dominant TSS positions (genomic coordinates with maximum TSS signal per consensus TSS cluster) of each group. Using the dominant TSS positions from two groups, the direction of the TSS shift was defined. If a promoter called for TSS shift has the same pre-and post-dominant TSS positions, this indicates that this promoter has the spatial changes in TSS signal distribution inside the promoter with the dominant TSS position remaining the same. For the heatmaps for TSS change upon each TF degradation, we used only the TSS that changed the dominant TSS position to characterize the effect of TSS shift magnitude (i.e. shift size).

#### Counting NET-CAGE reads

Read counts over 41 nt regions (±20) centered on TSSs were generated using the QuasR function qCount, with the option, orientation = “same” and without shifting (to quantify 5’ position of transcripts). For the gene expression change, fold change estimates were derived using the edgeR predFC function with a pseudocount of 8. For each differential analysis comparison, genes were considered to be differentially expressed with a false discovery rate (FDR) < 0.05 & an absolute log2 fold change > 0.5 to define moderate to strong gene expression response. For the comparison between transcriptional effect on TSS (41 nt windows as above) versus promoter-wide, promoter-wide effects were defined by the NET-CAGE signal change at 1,001 nt regions (-800/+200) relative to TSSs.

Meta profiles (heatmap) for NET-CAGE signal changes were generated using the QuasR function qProfile with the option orientation = “same” and normalized by scaling the total mapped reads to 30e6 reads. Profiles were then smoothed with a running mean of 51 nt and the mean signal of replicates is shown.

### Visualization of genome-wide coverage tracks

Integrative genomics viewer (v2.16.0) ^121^ was employed for visualization. Genome-wide coverage tracks (bigwig) were generated with bamCoverage ^122^ using 1 bp binsize (-bs), 51 bp smoothing (--smoothLength), the ENCODE blacklisted regions exclusion (--blackListFileName), CPM normalization (--normalizeUsing), normalization with chr 1 to 19, X, Y (--ignoreForNormalization) for ChIP-seq and RNA-seq. For ATAC-seq, the same parameters were used adding the options --Offset 1 and --extendReads 0. For NET-CAGE data, we used a publicly available pipeline (https://github.com/suimye/cage_tutorial/blob/master/cage.counting.pipeline.b0.01.s <u>h</u>). Here, mapped reads (only read1) were filtered to retain high-quality primary alignments (a mapping-quality ≥ 20) using samtools. Strand-specific CAGE TSS profiles were then generated with BEDtools genomeCoverageBed, counting only the 5′ ends of reads (-5) separately for the forward and reverse strands. The resulting bedGraph files were sorted and converted to BigWig format using UCSC bedGraphToBigWig, producing strand-resolved CAGE TSS signal tracks suitable for genome browser visualization. For each gene shown, the genomic tracks are oriented with the 5′ end on the left and the 3′ end on the right. NET-CAGE signal at promoter regions is represented using IGV tracks that show reads originating from the same strand as their respective genes, unless otherwise specified to display both strands.

### Locus-specific CAGE

Locus-specific CAGE reads were pre-selected to remove PCR-duplicated reads using UMIs. Using the R package ShortRead function readFastq ^112^, the reads that correctly contained library-specific sequences were selected (TGCATATCAACGCAGAGTACG at the 1^st^-21^st^ nucleotides of Read1, 5’ linker sequences ligated to the 5’ of transcripts). The selected reads were then assigned to each promoter using barcoded DNA (the 37^th^-52^nd^ nucleotides of Read2 CCTAGGN(10), N(10) is a promoter specific barcode). If multiple reads from the same promoter contained the same UMIs (the 1^st^-12^th^ nucleotides of Read2), they were considered to be PCR duplicates and only one read was selected. These de-duplicated reads were trimmed by removing the 1^st^-21^st^ nucleotides of Read1 and the 1^st^-12^th^ nucleotides of Read2 and mapped on the insert sequences using the QuasR function qAlign as in RNA-seq. The sequences of the inserts and barcodes are listed in Supplementary Table 4. The mapped reads were then used to quantify TSS signal by counting the first base of the mapped reads using the publicly available pipeline (https://github.com/suimye/cage_tutorial/blob/master/cage.counting.pipeline.b0.01.s <u>h</u>) as we did for NET-CAGE data. To normalize the TSS signal from the integrated promoters by the RMCE insertion ratio in pools, we estimated the insertion ratio from the barcoded DNA sequencing. The reads that contain universal sequences of the insert amplicons (CTGTGGTTGGTGTTCAGTCCCCTAGG at the 1^st^-26^th^ nucleotides of Read2) were selected using the ShortRead function readFastq and the promoter-unique barcodes (the 27^th^-36^th^ nucleotides of Read2) were counted. The ratio of the inserts was used to normalize the TSS signal of each promoter by dividing the TSS signal value by the normalized genomic DNA counts, representing the relative values for TSS signal per cell number as previously described for a similar assay ^42^. The mean of the normalized values for TSS signal was used for visualization of locus-specific CAGE signal. The barcodes for promoters are listed together with the construct designs in Supplementary Table 4.

### Single-molecule footprinting analysis

We analyzed the single-molecule footprinting data as previously described ^65,90^. Reads were first trimmed using Trimmomatic version 0.32 ^123^ using a paired end mode with the options (ILLUMINACLIP:TruSeq3-PE.fa:2:30:10, SLIDINGWINDOW:4:15, MINLEN:50).

The resulting adaptor-trimmed reads were mapped on inserted constructs using QuasR using a mode for bisulfite-converted templates (bisulfite = “undir”) with the option (alignmentParameter =-e 300-X 800-k 2 --best --strata). Methylation of Cs in the CpG and GpC contexts was then quantified using QuasR function qMeth and Cs with coverage <10 were excluded.

To distinguish endogenous CpG methylation and exogenous GpC methylation on GCG, we quantified endogenous CpG methylation levels on GCG using the samples without GpC methyltransferase treatment and only included GCG that showed low CpG methylation level (<20%). The resulting GpC methylation level was shown as 100 - methylation (%) in all figures. The promoter sequences used for analysis are listed in Supplementary Table 4.

## Acknowledgements

We thank members of the Schübeler laboratory, in particular, Ana Petracovici, Lisa Baumgartner, Julio Liu and Maria Eleni Vathi for their critical feedback on this study and Marc Timmers and Luca Giorgetti for input on the manuscript. We thank the functional genomics platform of the FMI especially Sirisha Aluri and Eliza Pandini Figueiredo Moreno for next-generation sequencing support, Hubertus Kohler for cell sorting, Panagiotis Papasaikas for computational support, Gergely Tihanyi for input on barcoded DNA sequencing for locus-specific CAGE, Arjun Udupa for technical support for cell sorting.

F. Moribe acknowledges support from the Nakajima Foundation and the Murata Overseas Scholarship Foundation. D. Schübeler acknowledges support from the Novartis Research Foundation, the Swiss National Science Foundation (310030_212716) and the European Research Council (EU Horizon 2020 ERC Advanced Grant; 884664).

## Author Contributions

F.M. and D.S. conceived and designed experiments. F.M. conducted cell line generation, genomics experiments and performed computational analysis. M.I. developed and implemented the CNN analysis and performed bioinformatics analysis.

S.D. provided mentorship and guidance in experiments and analysis and performed the quantitative imaging. C.W. performed western blotting experiments. L.H. assisted with genomics experiments and conducted NRF1 cell line generation and qPCR measurements. S.A.S. advised on the protocol for locus-specific CAGE and oversaw the next-generation-sequencing data. L.B. supervised computational analysis. D.S. supervised the project. F.M. and D.S. wrote the manuscript with input from the co-authors.

## Supplementary information

Supplemental Tables

S1: Constructs and gRNA sequences for cell line generation

S2: Antibody for western blotting and ChIP-seq

S3: NET-CAGE library preparation

S4: Sequences of promoter variant constructs for locus-specific CAGE

S5: Locus-specific CAGE library preparation

S6: Single-molecule footprinting library preparation

S7: PCR primers for distal TSS reporter

## Extended Data Figures

**Extended Data Fig. 1.**
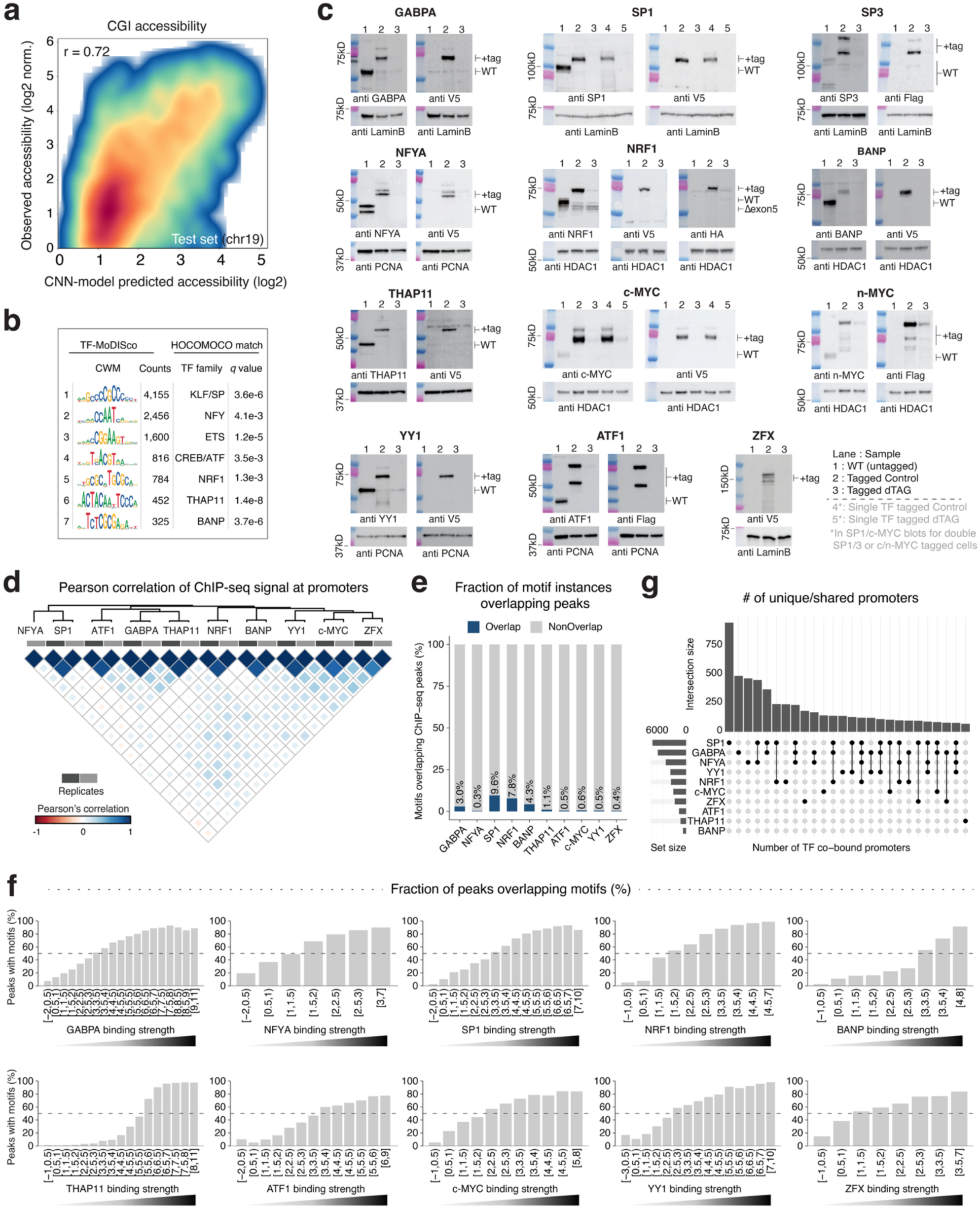
**a,** Comparison of chromatin accessibility (log2) at CGIs predicted by the CNN model and experimentally observed (r: Pearson’s correlation). **b,** Top de novo motif seqlets predicted to contribute to increased accessibility at CGIs. Known motifs in HOCOMOCO are assigned. **c,** Western blotting of degron-tagged TFs (500 nM dTAG13 for 4 hours). Each TF is tested with antibodies against the endogenous protein peptide and epitope tag. BANP degron cells were generated in our previous study ^45^**. d,** Pairwise correlation among ChIP-seq datasets, with color indicating Pearson’s correlation of ChIP-seq enrichment across all promoters (n = 24,575). **e,** Fraction of genome-wide TF motifs overlapping with ChIP-seq peaks. **f,** Fraction of ChIP-seq peaks containing the respective TF motifs, grouped by binding strength at peaks (ChIP-seq log2 enrichment). **g,** Counts of uniquely-bound or shared promoters at all promoters. Top: counts of promoters bound by either a single TF or multiple TFs, (pairs indicated by dotted lines). Left: total counts of promoters bound by each TF.

**Extended Data Fig. 2.**
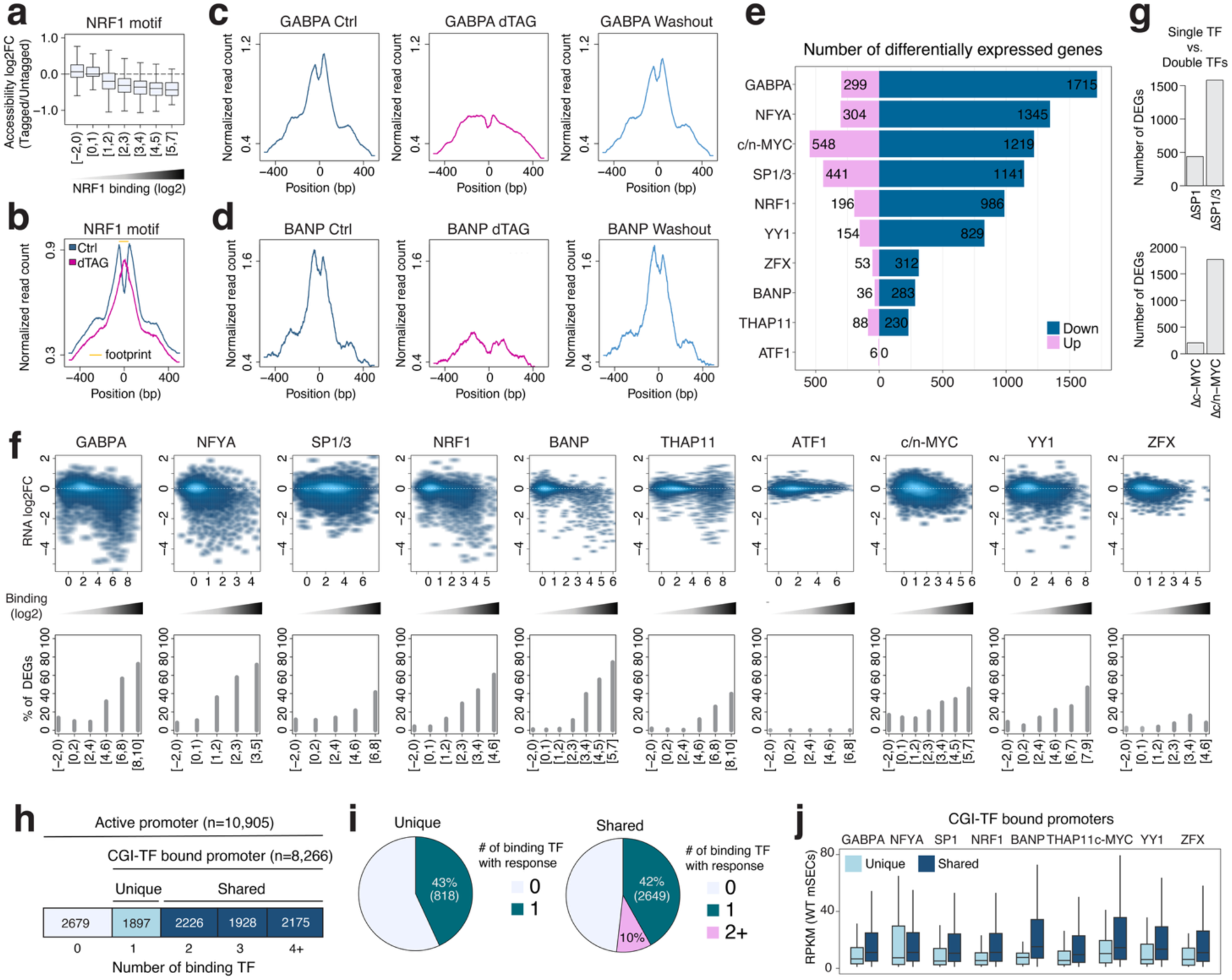
**a,** Reduction in accessibility upon NRF1 tagging. ATAC-seq log2 fold change between NRF1-tagged (non-degraded) and WT (untagged) cells at NRF1 motifs grouped by NRF1 binding strength; bin labels indicate ChIP-seq log2 enrichment of the bin. **b,** Accessibility change upon NRF1 degradation at the top 1,000 NRF1 binding sites (500 nM dTAG13 for 4 hours). Reduced ATAC-seq signal around the motifs is accompanied by NRF1 footprint loss centered on the motifs, locally increasing accessibility. **c, d,** Accessibility recovery after dTAG washout. ATAC-seq signal around TF motifs under control, dTAG and washout (dTAG for 4 hours followed by 16 hours recovery) conditions. Motifs with significant accessibility change upon dTAG treatment are analyzed (FDR < 0.05 & |log2 fold change| > 0.5, GABPA: n = 891, BANP: n = 99). **e,** Differentially expressed gene counts (DEG; FDR < 0.05 & |log2 fold change| > 0.5) upon TF degradation. **f,** Transcriptional responses of active genes (n = 10,095) upon TF degradation compared to binding strength at promoters. Top: ChIP-seq enrichment (log2) versus RNA-seq fold change (log2, dTAG/Ctrl). Bottom: Fraction of DEGs across promoters grouped by binding strength; bin labels indicate ChIP-seq enrichment of the bin. **g,** DEG counts upon dTAG treatment in single TF-tagged versus double TF-tagged cell lines. **h,** Promoter counts bound by different numbers of TFs (0/1/2/3/4+). **i,** Uniquely bound (n=1,897) or shared (n=6,329) promoters grouped by the number of TFs (0/1/2+) whose degradation causes transcriptional downregulation (FDR < 0.05 & log2 fold change <-0.5). **j,** Gene expression in WT mESCs comparing promoters bound by a single TF (Unique) or multiple TFs (Shared). Promoter counts (Unique/Shared) are GABPA: 384/3,915, NFYA: 218/1,969, SP1/3: 700/4,162, NRF1: 154/2,094, BANP: 25/450, THAP11: 62/790, c/n-MYC: 102/1,857, ZFX: 106/1,548.

**Extended Data Fig. 3.**
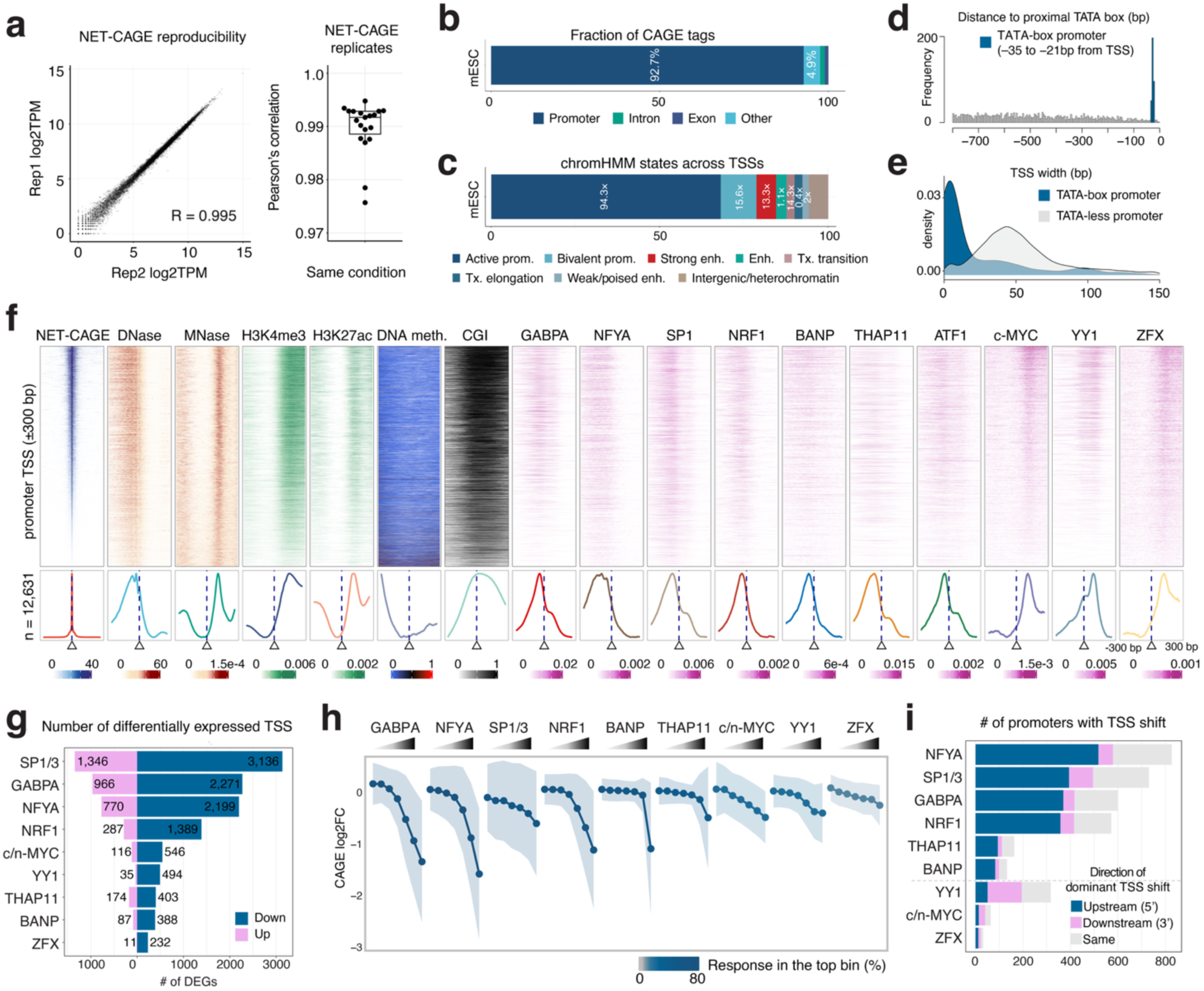
**a,** NET-CAGE reproducibility across replicates. Left: comparison of TSS signal (log2 TPM) in WT mESCs. Right: Pearson’s correlation between replicates across datasets (n=2). **b,** Genomic annotation for NET-CAGE TSSs in mESCs. **c,** ChromHMM states at NET-CAGE TSSs in mESCs; numbers indicate enrichment of each state. **d,** TATA-box positioning at promoters. TATA-box promoters contain a TATA motif-35 to-21 bp upstream of the TSS. Histograms show distance distribution between each TSS and the nearest upstream TATA box at active promoters. **e,** TSS width distribution for TATA-box (n = 346, median 7 bp) and TATA-less (n = 12,285, median 52 bp) promoters. **f,** TF binding sites (ChIP-seq) relative to TSSs at promoters (n = 12,631) with epigenome profiles. **g,** Differentially expressed TSS counts (FDR < 0.05 & |log2 fold change| > 0.5) upon TF degradation (500 nM dTAG13 for 4 hours). **h,** Changes in NET-CAGE signal upon TF degradation as a function of TF binding strength at promoters. Mean NET-CAGE fold change (log2) at active promoters (n = 12,631) grouped by binding strength of the respective TF (ChIP-seq enrichment ranked by 20-20-20-15-10-10-5%, from weakest to strongest). Shaded areas represent standard deviation, and line colors represent fraction of significant response in the top bin (FDR < 0.05 & |log2 fold change| > 0.5). **i,** TSS shift counts with color indicating shift direction (no change, upstream, downstream), defined by the relative position of dominant TSSs before and after dTAG treatment (see **Methods**).

**Extended Data Fig. 4.**
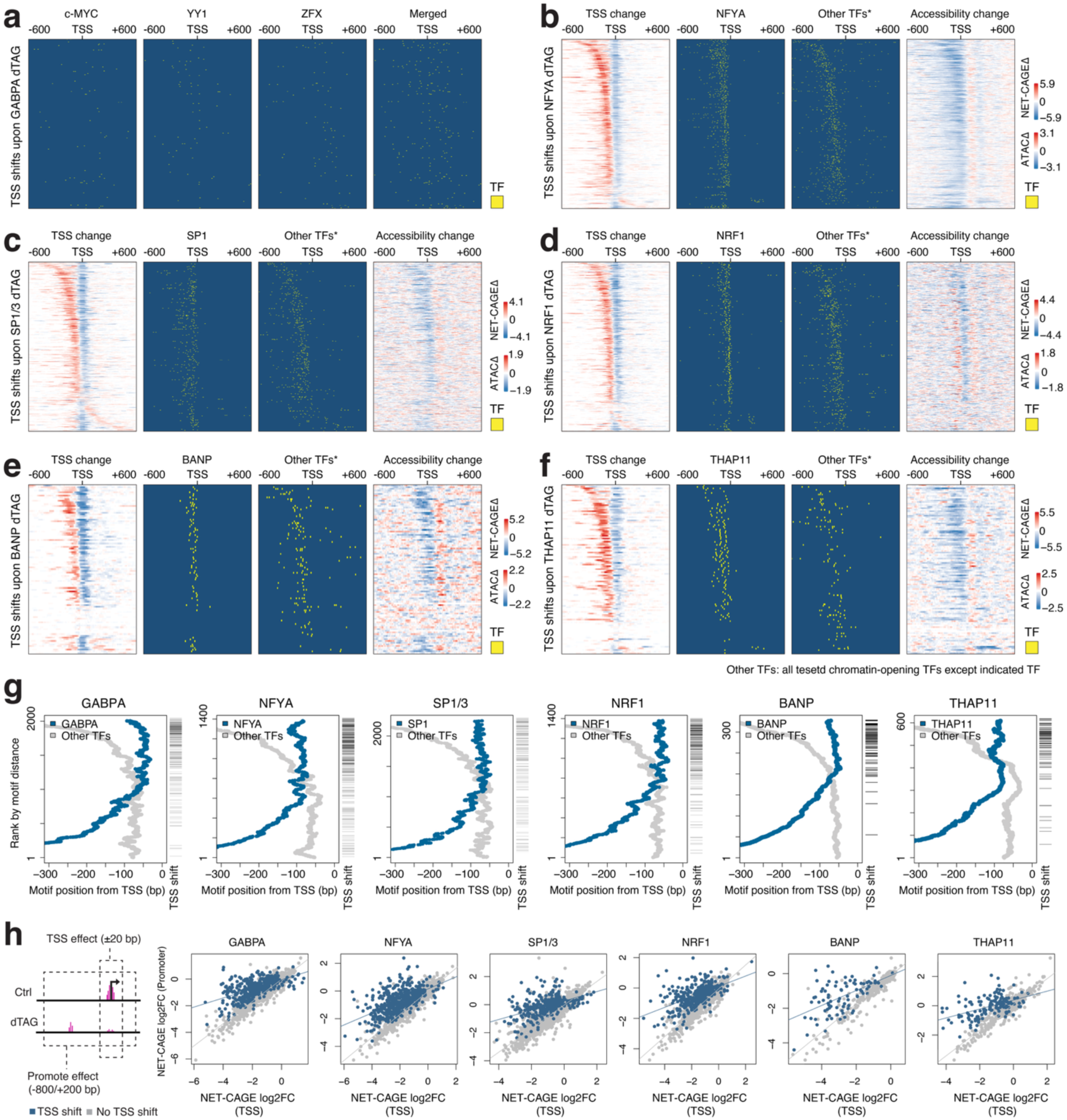
**a,** Positions of non-chromatin-opening TFs at promoters with TSS shifts upon GABPA degradation (n = 417). YY1, c-MYC and ZFX do not align with the shifted TSS positions. **b-f,** TSS shifts following degradation of chromatin-opening TFs. Promoters inducing a TSS shift upon each TF degradation are shown. Panels show NET-CAGE signal fold change (log2, dTAG/control), bound motifs of the degraded TFs, bound motifs of other chromatin-opening TFs (=*Other TFs), and ATAC-seq signal log2 fold change (dTAG/control). Promoter counts with TSS shifts shown are 579 in **(b)**, 495 in **(c)**, 415 in **(d)**, 99 in **(e)** and 111 in **(f)**. **g,** TSS shifts depend on the position of chromatin-opening TF. The occurrence of TSS shifts is shown in bars sorted by the distance between the degraded chromatin-opening TF (blue) and another proximal chromatin-opening TF (gray) at shared promoters. Colored lines represent average bound-motif positions, computed with a sliding 51-promoter window. **h,** Transcriptional impact of TSS shifts at TSSs and promoters. TSS effects are defined by NET-CAGE fold change (log2, dTAG/Ctrl) over ± 20 bp relative to TSSs and promoter effects are defined by NET-CAGE change over-800/+200 bp relative to TSSs. The lines indicate linear regression of each group.

**Extended Data Fig. 5.**
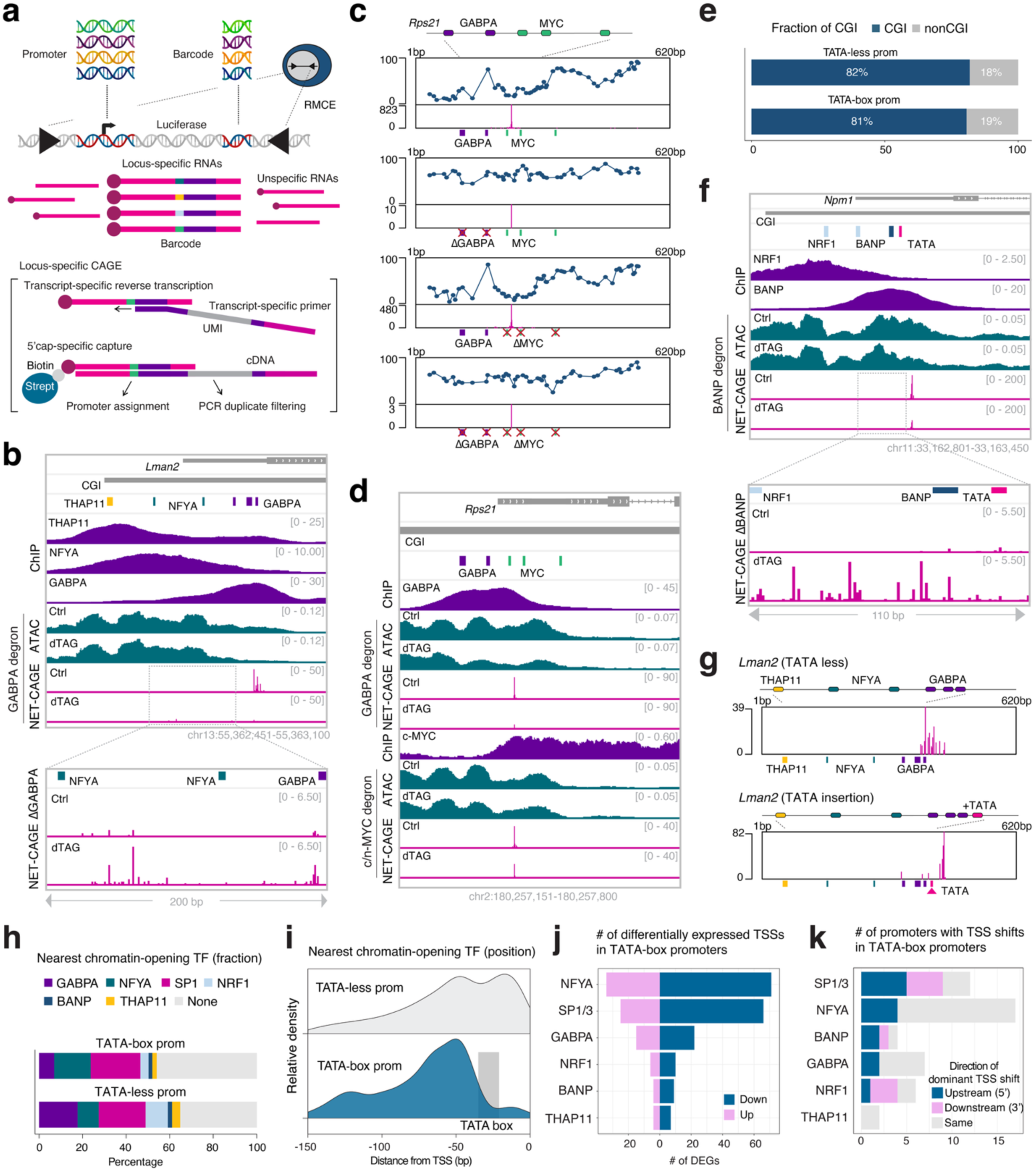
**a,** Schematic of locus-specific CAGE. **b,** Endogenous *Lman2* locus. Chromatin accessibility and TSSs at the endogenous promoter are recapitulated by the integrated *Lman2* promoters (**Fig. 4b**). **c,** GABPA-MYC containing promoter (*Rps21*). TSS signal and SMF readouts measured with/without GABPA/MYC motifs. **d,** Endogenous *Rps21* locus mirrored by the integrated *Rps21* promoters. **e,** Fraction of TATA-box and TATA-less promoters (n = 346 and n = 12,285, respectively) overlapping with CGIs. **f,** Representative TATA-box-containing CGI promoter (*Npm1*) showing a TSS shift upon BANP degradation. **g,** TATA box insertion into a CGI promoter (*Lman2*) converts a broad TSS profile into a sharp profile. **h,** Fraction of TATA-box and TATA-less promoters with chromatin-opening TF binding upstream of the TSS. The nearest TSS-proximal chromatin-opening TF (-800 to 0 bp) is indicated. **i,** Positional distributions of chromatin-opening TFs upstream of the TSS in TATA-box and TATA-less promoters. **j,** Differentially expressed TSS counts upon TF degradation for TATA-box promoters (FDR < 0.05 & |log2 fold change| > 0.5). 58% show differential expression. **k,** TSS shift counts upon TF degradation at TATA-box promoters with color indicating shift direction (no change, upstream, downstream), defined by the relative position of dominant TSSs before and after dTAG treatment. Compared to TATA-less promoters, more promoters retain the dominant TSS position, likely because TATA-box promoters have particularly strong dominant TSSs before dTAG treatment.

**Extended Data Fig. 6.**
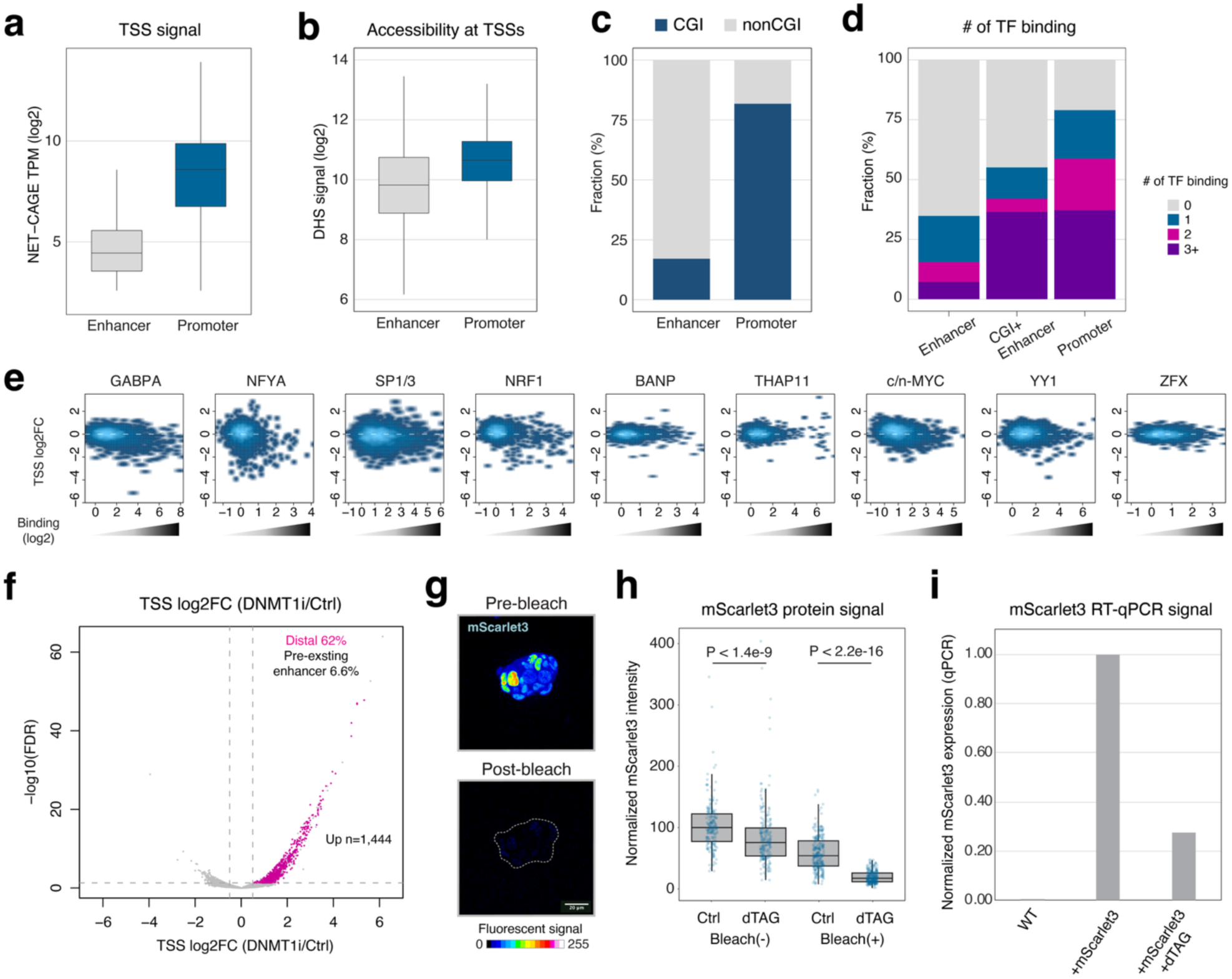
**a,**Comparison of TSS activity (NET-CAGE signal) between promoters (n = 12,631) and candidate enhancers (n = 2,640). **b,** Similar to **(a)** but for chromatin accessibility (DNase-seq signal) **c,** Fraction of promoters and candidate enhancers overlapping CGIs. **d,** Number of binding TFs at CGI promoters versus CGI enhancers. **e,** TSS changes at candidate enhancers upon TF degradation compared to binding strength at enhancers. X-axis: ChIP-seq enrichment (log2); Y-axis: NET-CAGE fold change (log2, dTAG/Ctrl). **f,** TSS expression changes after DNA methylation removal using a DNMT1 inhibitor (500 nM GSK-3484862 for 72 hours). Among the upregulated TSSs (FDR < 0.05 & log2 fold change > 0.5), the distal TSSs are colored in magenta. **g,** Representative microscopic images of mScarlet3 protein before and after bleaching. **h,** Imaging-based quantification of mScarlet3 protein upon GABPA degradation (500 nM dTAG13 for 12 hours). Wilcoxon rank-sum test: dTAG (n = 201) versus Ctrl (n = 197), *P*-value = 2.2e-16; and bleached dTAG (n = 254) versus bleached Ctrl (n = 264), *P*-value = 1.4e-09. **i,** RT-qPCR quantification of mScarlet3 transcript levels upon GABPA degradation (500 nM dTAG13 for 8 hours).

